# Divergence in alternative polyadenylation contributes to gene regulatory differences between humans and chimpanzees

**DOI:** 10.1101/2020.08.27.270686

**Authors:** Briana Mittleman, Sebastian Pott, Shane Warland, Kenneth Barr, Claudia Cuevas, Yoav Gilad

**Affiliations:** Genetics, Genomics and Systems Biology, University of Chicago, IL; Department of Human Genetics, University of Chicago, IL; Section of Genetic Medicine, Department of Medicine, University of Chicago, IL

**Author notes:** **Correspondence should be addressed to Y.G**.

## Abstract

Comparative functional genomic studies have shown that differences in gene expression between species can often be explained by corresponding inter-species differences in genetic and epigenetic regulatory mechanisms. In the quest to understand gene regulatory evolution in primates, the role of co-transcriptional regulatory mechanisms, such as alternative polyadenylation (**APA**), have so far received little attention. To begin addressing this gap, we studied APA in lymphoblastoid cell lines from six humans and six chimpanzees, and estimated usage for 44,432 polyadenylation sites (**PAS**) in 9,518 genes in both species. While APA is largely conserved in humans and chimpanzees, we identified 1,705 genes with significantly different PAS usage (FDR of 0.05) between the two species. We found that genes with divergent APA patterns are enriched among differentially expressed genes, as well as among genes that show differences in protein translation between species. In particular, differences in APA between humans and chimpanzees can explain a subset of observed inter-species protein expression differences that do not display corresponding differences at the transcript level. Finally, we focused on genes that have a dominant PAS, namely a PAS that is used more often than all others. Dominant PAS are highly conserved, and inter-species differences in dominant PAS are particularly enriched for genes that also show expression differences between the species. This study establishes APA as another key mechanism underlying the genetic regulation of transcript and protein expression levels in primates.

## Introduction

Humans and our close primate relatives exhibit a striking array of phenotypic diversity despite sharing homologous proteins with nearly identical amino acid sequences. Understanding how this diversity is propagated from genomic sequence to mRNA and then to protein necessitates an understanding of the regulatory mechanisms that occur before, during, and after transcription. Studying gene regulatory features in humans and other primates has long provided opportunities to understand genome evolution and function. For example, studies comparing patterns of epigenetic marks in primates have provided mechanistic explanations that link genetic variation and divergence to differences in gene expression levels (Banovich et al., 2014; Cain et al., 2011; McVicker et al., 2013; Pai et al., 2011). Although many studies have focused on inter-species differences in the regulation of gene expression, fewer studies have addressed isoform-level variation, which contributes to differences in mRNA, translation, and protein levels between species.

The main mechanisms that contribute to mRNA isoform diversity are alternative splicing and alternative polyadenylation (**APA**). Alternative splicing produces different combinations of coding sequences in mature mRNA and protein. APA occurs at genes that have more than one polyadenylation site (**PAS**) and can result in mRNAs with different coding sequences or variable 3’UTR lengths. Like alternative splicing, APA that occurs within the gene body can affect protein sequence and function (Lee et al., 2018; Pan et al., 2006; Sandberg et al., 2008; Tian & Manley, 2017, p. 20; Vasudevan et al., 2002; Yao et al., 2018). APA that occurs outside of the coding sequence, in the 3’UTRs, can lead to differential inclusion of protein-binding motifs that can affect translational efficiency, mRNA stability, and mRNA localization (Mayr, 2017; Tian & Manley, 2017). Yet, despite its potential to produce tremendous variation in mRNA and protein regulation, few studies have explored the contribution of APA to regulatory divergence between species. Indeed, our current understanding of APA conservation in mammals comes from few comparative studies of humans and rodents (Ara et al., 2006; R. Wang et al., 2018). However, these studies used sequence conservation rather than direct measurements of PAS usage to characterize APA (R. Wang et al., 2018). Thus, it remains possible that many mammalian PAS are functionally divergent despite having similar sequences.

To gain insight into APA conservation in humans and chimpanzees and understand how differences in APA contribute to gene regulation, we performed 3’ sequencing (**3’ Seq**) of mRNA isolated from nuclei collected from human and chimpanzee LCLs. We integrated PAS usage measurements with RNA-sequencing data collected from the same cell lines to understand the relationship between APA and gene expression levels. Finally, we used ribosome profiling and protein measurements previously collected in the same panel of human and chimpanzee LCLs to explore the effects of APA on protein levels (Khan et al., 2013; S. H. Wang et al., 2018), reasoning that an understanding of how APA isoform usage varies among primates could help explain why some human and chimpanzee genes are differentially expressed at either the mRNA or protein levels, but not both.

## Results

### Describing alternative polyadenylation in human and chimpanzee LCLs

We performed 3’ Seq of mRNA from 6 human and 6 chimpanzee lymphoblastoid cell lines (**LCLs**), which we have previously used to study a variety of other functional genomic phenotypes (Cain et al., 2011; Khan et al., 2013; S. H. Wang et al., 2018; Zhou et al., 2014). We collected mRNA separately from whole cells and isolated nuclei. The two cellular fractions serve as biological replicates, which we mainly used to examine the quality of our data (**methods**). By collecting data from isolated nuclei, we were able to capture polyadenylated transcripts before they became undetectable due to other regulatory processes, such as isoform-specific decay (Mittleman et al., 2020).

We mapped human 3’ Seq reads to the GRCh38 reference genome and chimpanzee 3’ Seq reads to the panTro6 reference genome (Chimpanzee Sequencing and Analysis Consortium, 2005; Schneider et al., 2017) (**methods**). 3’ Seq relies on a poly(dT) primer to target the poly(A) tail of mRNA molecules; however, it can also mis-prime by binding a sequence of genomic adenines. To account for mis-priming of off-target genomic sequences we removed reads that mapped to genomic regions containing ≥70% adenine or 6 consecutive adenine bases in the 10 bp directly upstream of the mapped location (Mittleman et al., 2020; Sheppard et al., 2013; Tian et al., 2005) (**methods**). In addition, we treated all ambiguous nucleotide positions as adenines to ensure that differences in reference genome quality did not bias the detection of polyadenylation sites (**PAS**) or mis-priming events **(methods)**. As expected, the filtered aligned sequences, in both species, were enriched at transcription end sites (**TES**) and showed a similar distribution along orthologous 3’ UTRs **(methods, Figure 1 - figure supplement 1**). Next, we used a custom peak calling method to ascertain PAS in humans and chimpanzees separately (**methods**).

**Figure 1:**
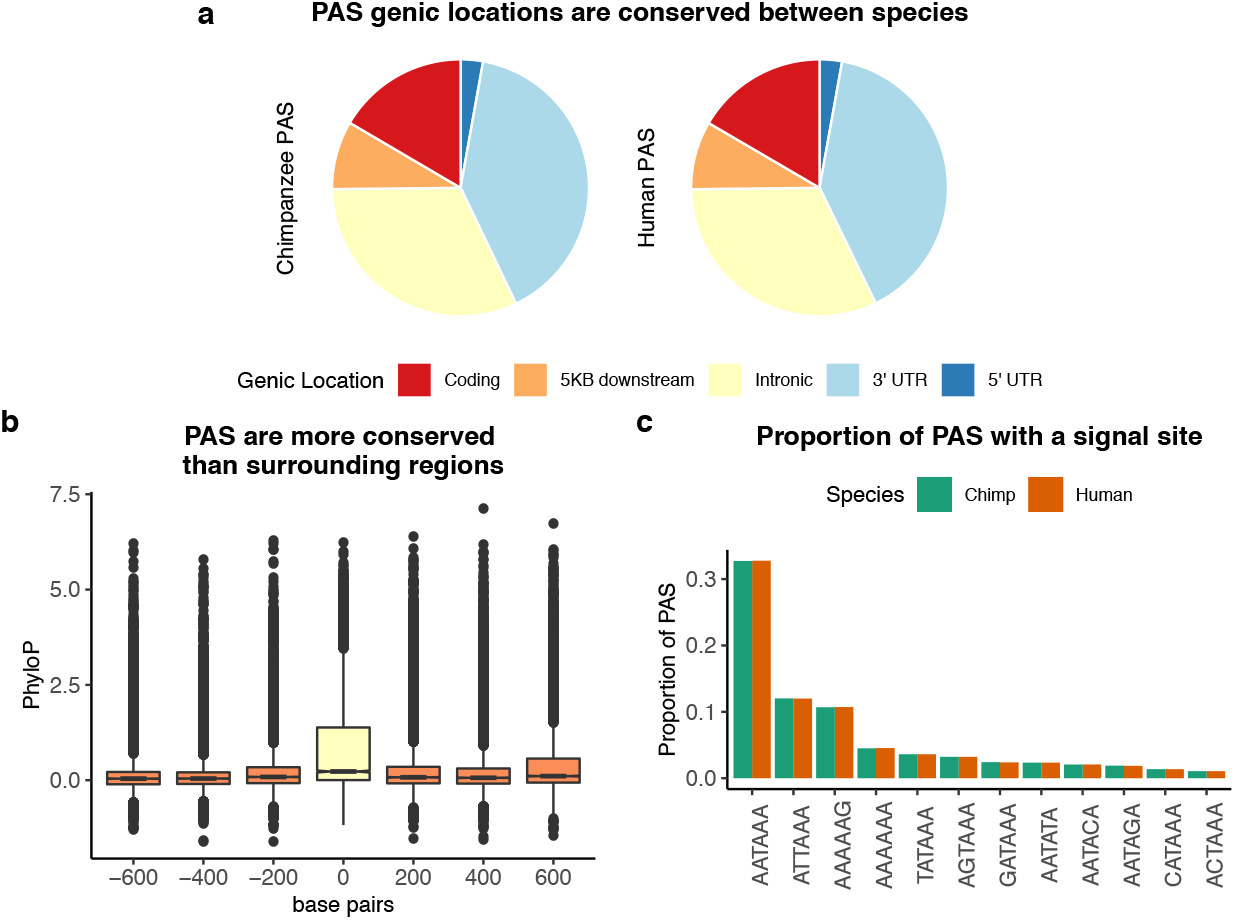
Sequence conservation of PAS between humans and chimpanzees. a. Genic locations for 44,074 PAS identified in chimpanzee (left) and 44,130 PAS identified in human (right). b. Mean PhyloP scores for PAS regions (yellow) as well as three 200bp regions upstream and downstream (orange). c. Proportion of human and chimpanzee PAS regions with each of the 12 annotated signal site motifs from Beaudoing et al (Beaudoing et al., 2000).

To compare PAS usage across species, we needed to identify the orthologous genomic regions of all PAS in our dataset, regardless of the species in which they were originally annotated. As we were unable to confidently identify orthologous PAS at base pair resolution (inferring synteny at base pair resolution in non-coding regions is challenging; (Broad Institute Sequencing Platform and Whole Genome Assembly Team et al., 2011)), we extended each PAS by 100 bp upstream and downstream. We then used a reciprocal liftover pipeline to obtain an inclusive set of PAS regions with which we could confidently compare PAS usage between species (**methods**).

To quantify PAS usage, we first assigned each PAS to a gene using the hg38 RefSeq annotation (Pruitt et al., 2005). We then computed usage for each PAS in each individual as the fraction of reads mapping to one PAS over the total number of reads mapping to any PAS for the same gene (**Figure 1 - figure supplement 2**). We excluded PAS in lowly expressed genes or with less than 5% usage, as measurements from sparse data are highly susceptible to random error (**methods**). We observed a strong correlation between PAS usage in mRNA from the nuclear and total cell fractions in all but one cell line (human NA18499; **Figure 1 - figure Supplement 3**), which we subsequently excluded from the study. We re-identified PAS after removing all data from NA18499 and re-quantified PAS usage using nuclear 3’ Seq data from 5 human and 6 chimpanzee LCLs. Using this analysis pipeline we identified a total of 44,432 PAS in 9,518 genes, which we used for all downstream analyses. On a genome-wide scale, we found that mean PAS usage is highly correlated between species (Pearson’s correlation, 0.9, *p*<*2.2×10^−16^*, **Figure 1 - figure supplement 4)**. However, as expected, 41.8% of the variation in PAS usage (as explained by the top principal component of the data) is highly correlated with species (Pearson’s correlation 0.99, p =2.95×10^−8^, **Figure 1 - figure supplement 5**), indicating substantial divergence in PAS usage.

We used a number of analyses to confirm that our ability to detect PAS was not biased by gene expression level or species. If our ability to detect PAS were biased by gene expression, we might expect a positive correlation between gene expression level and the number of PAS we detected. In our data, the number of PAS per gene is negatively correlated with gene expression level in both species (**Figure 1 - figure supplement 6**, Human: Pearson’s correlation −0.17, *p* <*2.2×10^−16^*, Chimpanzee: Pearson’s correlation −0.19, *p* <*2.2×10^−16^*). If our ability to detect PAS were biased by species, we would expect to identify more PAS per gene in one species over the other. This is neither the case genome-wide nor when we test each gene independently. We identified, on average, 3.87 PAS per gene in humans and 3.46 PAS per gene in chimpanzees. On average, per gene, the number of PAS in human minus the number of PAS in chimpanzee is 0.39 with a median value of 0 (**Figure 1 - figure supplement 7**). Moreover, as expected, the physical distribution of PAS across genes is conserved, with the majority of PAS located in 3’ UTRs (17,688; 40% in chimpanzee and 17,620; 40% in human) and a considerable proportion located in introns (14,095; 32% in chimpanzee and 14,119; 32% in human) (**Figure 1A**).

To assess sequence conservation in PAS regions, we downloaded phyloP scores computed over 100 vertebrate genomes from the UCSC Genome Browser, and calculated mean phyloP scores in PAS regions. Higher mean phyloP scores correspond to regions of higher sequence conservation, and thus, slower evolution. (Pollard et al., 2010). Overall, sequence elements at PAS are more conserved than surrounding regions (**Figure 1B**, Wilcoxon rank sum test, *p*<*2.2×10^−16^*). This pattern also holds independently for PAS in all genic locations other than introns (**Figure 1 - figure supplement 8**).

We identified 302 and 357 human- and chimpanzee-specific PAS, respectively (**methods**). It has been previously shown that most PAS are directly preceded by one of 12 annotated sequence motifs that recruit cleavage and polyadenylation machinery to mRNA molecules as they are transcribed (Beaudoing et al., 2000) We asked if creation or disruption of a signal site motif could be responsible for species-specific PAS by mapping signal site motifs in both human and chimpanzee for each PAS region. Although human and chimpanzee PAS regions are equally likely to contain each of the 12 annotated signal sites (**Figure 1C**), only the top two most commonly used motifs, AATAAA and ATTAAA, are associated with increased PAS usage (**Figure 1 - figure supplement 9**). Thus, we considered only the presence or absence of these two motifs in subsequent analyses. Of the 302 human-specific PAS, 14 have human-specific signal sites and 6 have chimpanzee-specific signal sites. Of the 357 chimpanzee-specific PAS, 24 have a chimpanzee-specific signal site and 6 have a human-specific signal site. These numbers are small; still, species-specific signal sites are more abundant than expected by chance among species-specific PAS in human (5.7X, hypergeometric test, *p*=*2.30×10^−7^*) and in chimpanzee (8.3X, hypergeometric test, *p*=*3.2×10^−15^*), suggesting that signal site changes can explain a subset of differences in PAS usage. For example, we identified a chimpanzee-specific PAS about 1 kb upstream of a PAS used in both species in the 3’ UTR of *MAN2B2*. The ancestral signal site conserved in chimpanzee is AATAAA; however, there has been a T to C transition in the human lineage (Blanchette, 2004) (**Figure 1 - figure supplement 10**). This transition is likely responsible for the loss of PAS in humans.

### Characterizing inter-species differences in PAS usage

While a few hundred PAS are species-specific, the majority of PAS (98.5%) were identified in both species. We thus sought to characterize quantitative differences in alternative polyadenylation (**APA**) patterns between human and chimpanzee by estimating the difference in usage of individual PAS in each species. To do so, we used the leafcutter differential splicing tool (Y. I. Li et al., 2018), which allowed us to tests for differences in normalized PAS usage fractions while accounting for gene structure (**methods**). Using this approach, at an FDR of 5% we identified 2,342 PAS (in 1,705 genes) whose usage differs by 20% or more between the species (**Figure 2A**). We applied an arbitrary effect size cutoff to focus on larger inter-species differences, which are more likely to have functional consequences. The list of all PAS whose usage differs between the species, regardless of the effect size, is available in **Supplemental Table 1**.

**Figure 2:**
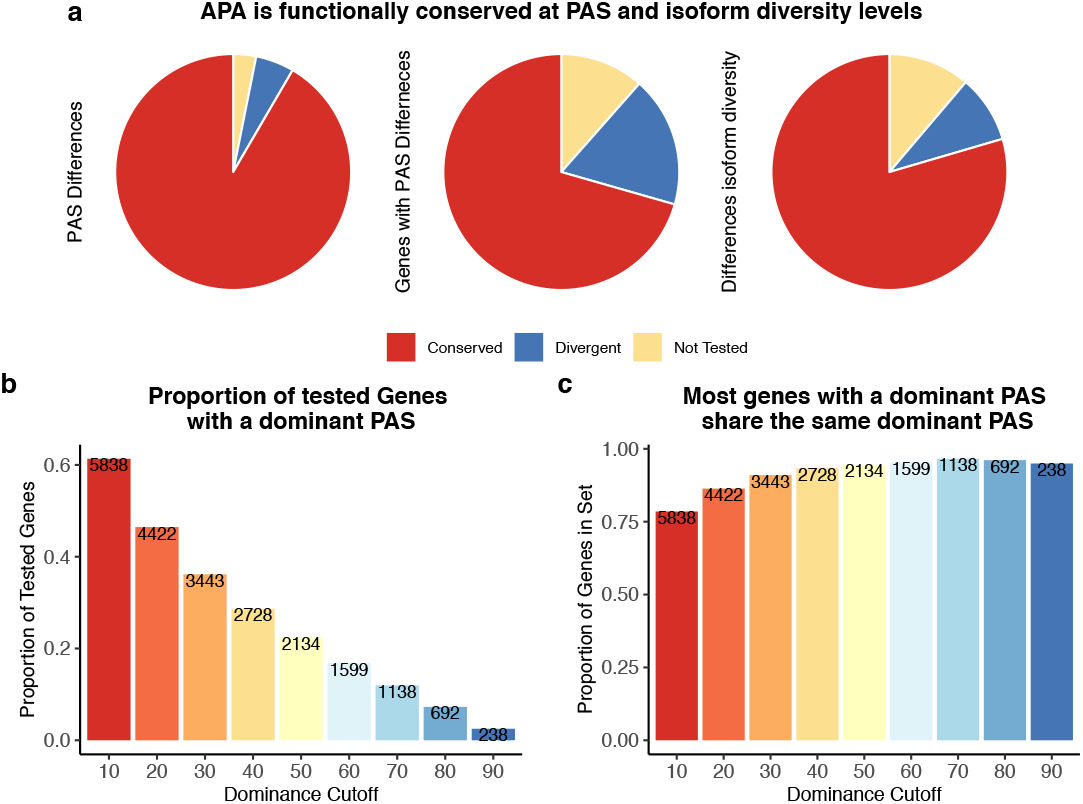
APA is functionally conserved in both species. a. Proportion of PAS and genes differentially used at PAS and isoform diversity level. (left) Divergent PAS are the 2,342 PAS differentially used at 5% FDR. Conserved are the PAS not differentially used at 5% FDR. Not tested PAS were removed from analysis by leafcutter tool. (middle) PAS level differentially used PAS reported at the gene level. Divergent genes are the 1,705 genes with PAS differentially used at 5% FDR. Conserved genes are the genes with no PAS differentially used at 5% FDR. (right) Divergent genes are the 881 genes with differences in isoform diversity between species at a 5% FDR. Conserved gene are genes without differences in isoform diversity. Genes with one PAS were not tested. b. Proportion of tested genes with a dominant PAS in either species according to a range of cutoffs. Number of genes are reported in bars. c. Proportion of the number of the number of genes with a dominant in either species that share the top used PAS according to each dominance cutoff. Number of genes with a dominant PAS is either species are reported in bars.

To better understand the mechanisms that underlie inter-species differences in PAS usage, and the potential functional impact of such differences, we considered the APA data in different contexts. First, we noticed that the spatial distribution of differentially used PAS reflects the distribution of all PAS; namely, differentially used PAS are most often located in 3’ UTRs, followed by introns (**Figure 2 - figure supplement 1**). Within the 3’ UTR, however, differentially used PAS are more frequently the first ones compared with PAS that are used similarly in the two species (**Figure 2 - figure supplement 2**, difference in proportion test, *p*=*0.0015*). This pattern is intriguing, because changes in the usage of the first PAS in the 3’ UTR may have the largest overall impact on the transcript length, and hence potentially the largest functional impact. However, it is also possible that we are more likely to detect differences in usage in the first PAS in the 3’UTR because this site is transcribed earlier, and our estimate of usage is relative to all other sites in each gene.

We therefore sought evidence that differences in PAS usage may have functional consequences. In a previous study, we identified genetic variants associated with variation in PAS usage (**apaQTLs**) in a panel of 52 human LCLs (Mittleman et al., 2020). We found that genes with inter-species differences in PAS usage are highly enriched for apaQTLs (160, empirical p-value based on 10,000 permutations *p*=*0.001*, **Figure 2 - figure supplement 3**). This observation indicates that inter-species differences in APA usage can often be found in genes whose regulation varies also at the population level, generally suggesting relaxation of evolutionary constraint on the regulation of such genes. We next considered sequence divergence at PAS by obtaining phyloP scores for all PAS flanking regions (200 bp, as explained above). If many changes in PAS usage are genetically controlled, we would expect genomic regions of differentially used PAS to be less conserved than regions containing PAS sites that have similar usage. Indeed, differentially used sites are enriched for regions with negative mean phyloP scores (1.02X, hypergeometric test, *p*=*0.02*). This observation indicates that sequence divergence is often associated with differences in PAS usage, and that the majority of PAS usage in humans and chimpanzees may be generally conserved due to evolutionary constraint.

We next asked, more specifically, if signal site changes are likely to lead to differences in PAS usage. We addressed this question by performing two analyses. First, we focused on the 82 differentially used PAS with a signal site that is annotated in only one of the species. We found that the presence of a species-specific signal site is associated with increased PAS usage, as might be expected (human enrichment 3.82X, hypergeometric *p*=*1.37×10^−10^*, chimpanzee enrichment 3.02X; p=3.91×10^−8^). Second, we considered the presence of G/U-rich elements, which are known signals to the molecular machinery for polyadenylation (Colgan & Manley, 1997). Specifically, we considered the proportion of uracil bases in the PAS regions. Despite a high correlation in overall uracil content in both species (Pearson’s correlation 0.99, *p*<*2.2×10^−16^*), the usage of PAS with greater uracil density in one species are more likely than expected by chance to be up-regulated in that species (Chimpanzee 1.04X enrichment; *p*=0.03, Human 1.06X enrichment; *p*=*0.03*). Though species-specific signal sites explain a modest proportion of inter-species differences in PAS usage, these cases demonstrate the link between sequence evolution and conservation of PAS usage.

### The relationship between differences in alternative polyadenylation and gene expression

Our analysis to this point indicates that inter-species differences in PAS usage are often genetically controlled, but generally we have not found strong evidence that they are functionally important. We explored this further by considering the APA data in the context of gene expression data that we collected from the same 6 human and 6 chimpanzee LCLs (see **methods** for data collection procedures and low-level analysis of the RNA-seq data). We found no meaningful correlation between inter-species differences in gene expression levels and changes in polyadenylation site usage (**ΔPAU**) in 7,462 genes for which we had both types of data (Pearson’s correlation = −0.06, p=3.1×10^−7^, **Figure 3A**). We then separately considered the data for the 3’ UTR and intronic PAS, because we previously found a different relationship between PAS usage in these genic regions and gene expression levels (Mittleman et al., 2020). Indeed, we found that inter-species differences in the usage of intronic and 3’ UTR PAS correlate with differences in expression effect size between the species at an equal magnitude, but in opposite directions (**Figure 3B**). Increased usage of intronic sites is correlated with increased expression levels, while increased usage of 3’ UTR sites is correlated with decreased expression.

**Figure 3:**
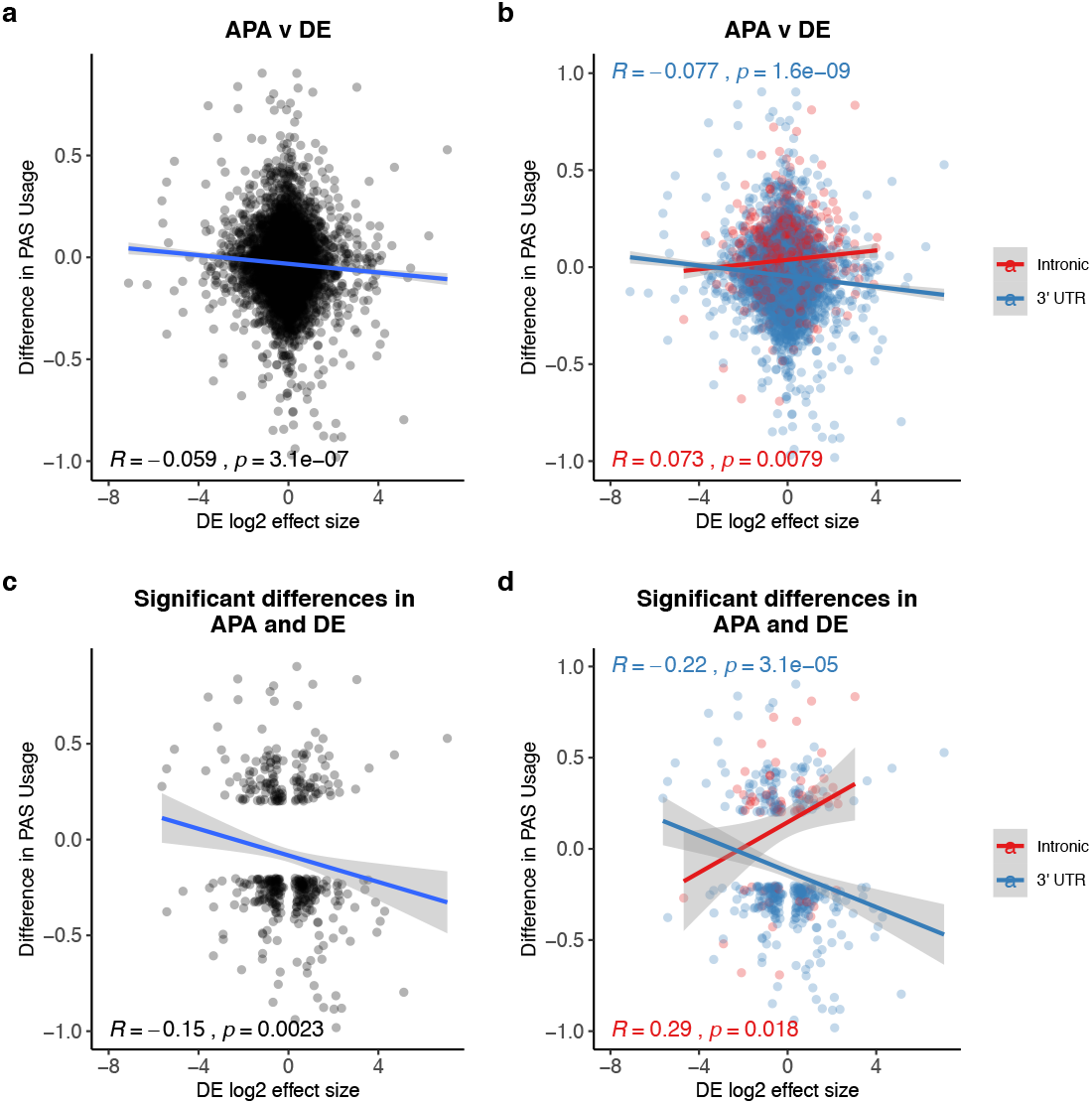
PAS usage differences for intronic and 3’ UTR PAS correlate with DE effect sizes at similar magnitudes but in opposite directions. a. ΔPAU for top intronic or 3’ UTR PAS per gene (methods) plotted against differential effect size from differential expression analysis. b. ΔPAU for top intronic or 3’ UTR PAS per gene (methods) plotted against differential effect size from differential expression analysis for genes with significant differences in each phenotype at 5% FDR. c. ΔPAU for top intronic or 3’ UTR PAS per gene (methods) plotted against differential effect size from differential expression analysis. d. ΔPAU for top intronic or 3’ UTR PAS per gene (methods) plotted against differential effect size from differential expression analysis for genes with significant differences in each phenotype at 5% FDR. In all panels, we calculated the linear regression and Pearson’s correlation. In all panels, negative ΔPAU and DE effect sizes represent upregulation in chimpanzees. In panels b and d, we colored the points and regressions by genic location.

We focused on 3,796 genes that were classified as differentially expressed between humans and chimpanzees at 5% FDR (**methods**). We found that genes with at least one differentially used PAS between the species are more likely to be classified as differentially expressed than expected by chance (610 genes, 1.12X enrichment, hypergeometric test, p=3.18×10^−5^). Examining the subset of 610 genes, we observed a modest but significant negative correlation between differential expression effect size and ΔPAU when we considered all PAS (Pearson’s correlation= −0.15, p=0.0023, **Figure 3C**). Separating the analysis by PAS genic location revealed, again, an opposite direction of the correlation between gene expression and the usage of either 3’ UTR or intronic PAS (**Figure 3D**). These observations are consistent when we use PAS data based on 3’ Seq data from whole cells instead of from the nuclear fractions, suggesting that the observed relationship is not due to nuclear export failure (**Figure 3 - figure supplement 1, methods**).

To provide possible mechanistic insight into the relationship between PAS usage and gene expression, we identified AU-rich elements in 3’ UTRs in both human and chimpanzee. AU-rich elements in 3’ UTRs have been linked to destabilization of mRNA transcripts and translation repression (Floor & Doudna, 2016; Moore et al., 2014; Siegel et al., 2020). We found that the 3’ UTRs of genes that show an inter-species difference in 3’UTR PAS usage have a higher number (Wilcoxon test, *p* < 10^−16^, Figure 3 – figure supplement 2) and density (*p* = 5.2×10^−6^, Figure 3 – figure supplement 2) of AU-rich elements compared with genes in which the 3’UTR PAS is similarly used in the two species.

### Considering overall APA diversity

We explored the relationship between inter-species differences in APA and gene expression by using a different perspective. We hypothesized that we could gain more insight into regulatory variation by summarizing the PAS diversity for a given gene using a single statistic, rather than by analyzing the usage of each site separately. To do so, we measured isoform diversity using Simpson’s D (**D**), a metric traditionally employed by ecologists to measure taxon diversity between environments (Morris et al., 2014). In our system, higher D values indicate that usage is spread more evenly across all PAS for a gene, while low D values suggest the one PAS is more dominant than others (**methods**). As expected, in both humans and chimpanzees, D values are correlated with the number of PAS per gene (**Figure 2 - figure supplement 4, 5, 6**; human Pearson’s correlation 0.62, p<2.2×10^−16^, chimpanzee Pearson’s correlation 0.63, p<2.2×10^−16^).

Using Simpson’s D values calculated for each gene in each individual, we identified (at 5% FDR) 881 genes with significant differences in isoform diversity between species (**Figure 2A, methods**). Of these, 426 are genes for which we did not previously detect an inter-species difference in PAS usage, indicating that Simpson’s D is capturing an additional dimension of, or is more sensitive to, APA variation between species (**Figure 2 - figure supplement 7, for example see Figure 5 - figure supplement 1**).

We proceeded by focusing on genes with low isoform diversity, suggesting a single dominant PAS. We calculated a dominance metric for each gene as the difference in mean usage between the first and second most used PAS (we used different cutoffs to classify dominance; see **methods**). We found that the classification dominant PAS is highly consistent across species; a result that is quite robust with respect to the approach used to classify PAS as dominant (**Figure 2B, 2C**). While the dominant PAS is the same for most genes in humans and chimpanzees, differences in usage of a dominant PAS are likely to contribute more to differential APA that have functional consequences between species than differences in other PAS. Indeed, regardless of the specific cutoff we used to define dominant PAS, when the dominant PAS is not the same in humans and chimpanzees, the corresponding genes are more likely to be differentially expressed between the species compared with genes where the dominant PAS is the same in both species, (for cutoffs between 0.2 and 0.7, all *p* < *0.005)*), and even compared with genes in which only a non-dominant PAS is differentially used (*p* > 0.8 for all cutoffs; **Figure 4A,B**).

**Figure 4:**
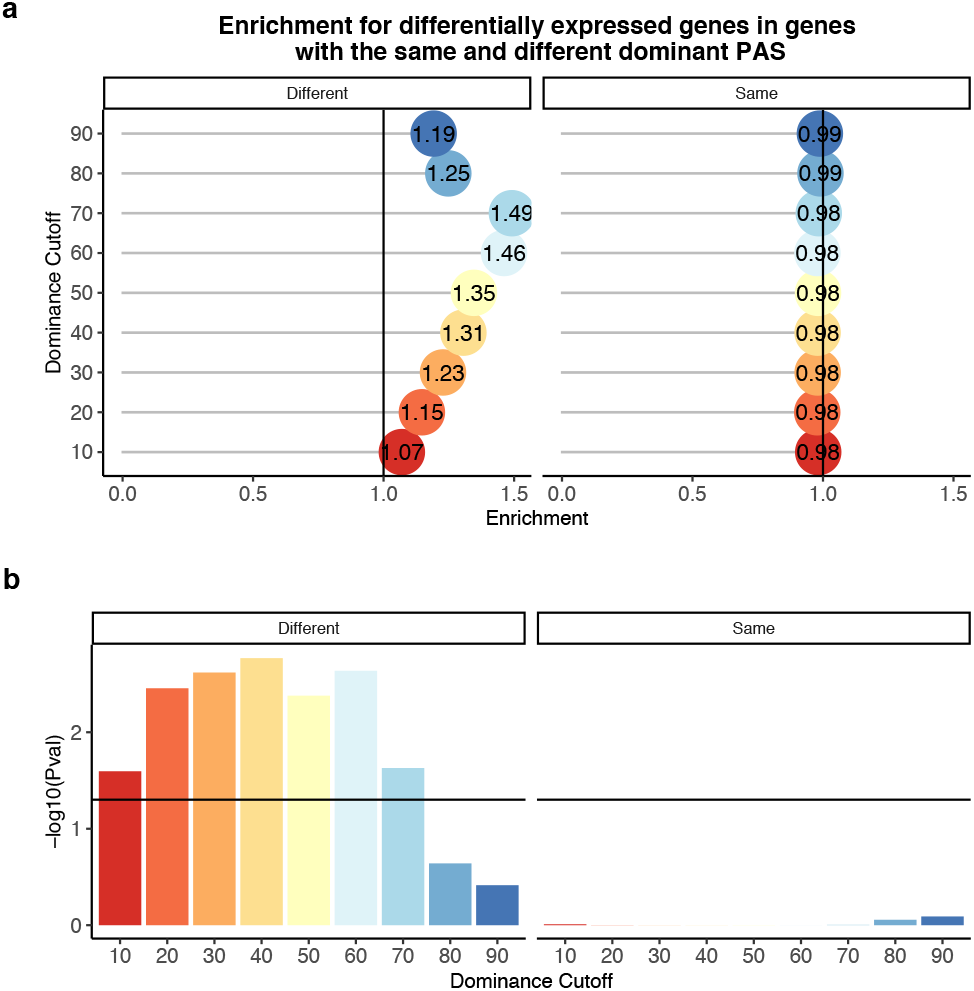
Difference in dominant PAS site between species likely drives differences in expression. a. Enrichment of genes with the different (left) of same (right) dominant PAS by dominant cutoff in differentially expressed genes. b. −log10(P-values) for enrichments in a calculated with hypergeometric tests. Horizontal line represents p-value of 0.05.

In a previous study that collected mRNA from a larger panel of human, chimpanzee, and rhesus macaque LCLs, Khan *et al.* identified genes whose regulation likely evolves under directional selection in humans and chimpanzees (Khan et al., 2013). We were able to consider RNA and protein expression data as well as APA data from 2,532 genes. We found that twenty-two of the genes with significant inter-species differences in APA at both the site level and in isoform diversity are among those whose regulation likely evolves under directional selection in the chimpanzee lineage, a 1.6X enrichment over what is expected by chance (hypergeometric test, p=0.015). We did not find a similar enrichment when we considered genes whose regulation evolved under selection in humans, but the sample size is rather small.

### Variation in APA and differences in protein expression

Given the well-characterized molecular connection between APA and the regulation of protein translation, we hypothesized that genes with inter-species differences in APA are also more likely to be differentially translated between the species (Di Giammartino et al., 2011; Floor & Doudna, 2016; Tian & Manley, 2017). To examine this, we obtained estimates of protein translation based on ribosome profiling data that were collected from human and chimpanzee LCLs by Wang *et al*. (S. H. Wang et al., 2018). At a 5% FDR, Wang *et al.* identified 1,993 differentially translated genes between humans and chimpanzees. Genes with significant inter-species differences in isoform diversity, but without significant differences in usage at individual PAS, are enriched among the differentially translated gene set (1.21X, hypergeometric test, P = 0.011; **Figure 5A,B**).

**Figure 5:**
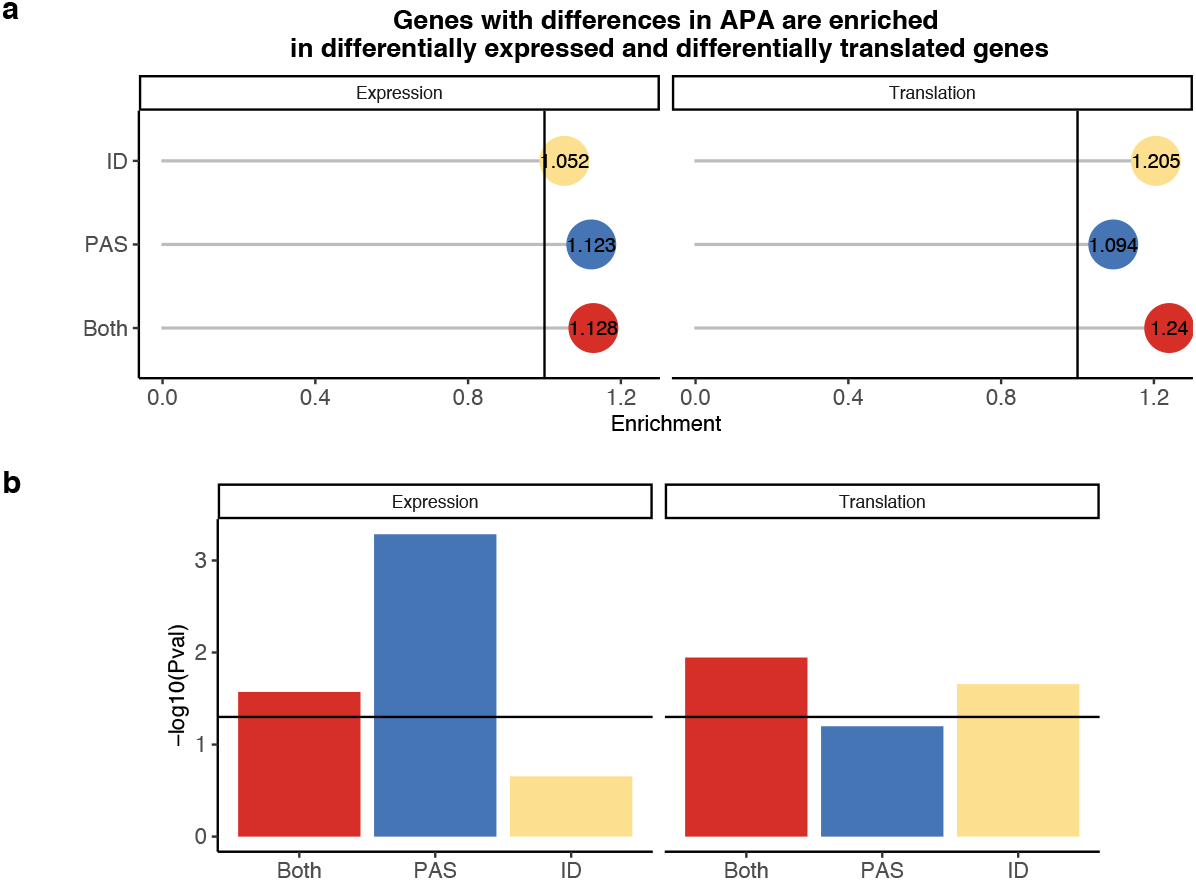
PAS level differences in APA may drive differences in expression while isoform diversity differences likely drive translation differences. a. Enrichment of genes with isoform level differences (ID), differences in APA at PAS level (PAS), or at both levels (Both) within differential expressed genes and differentially translated genes. Differentially translated genes reported by Wang et al. (S. H. Wang et al., 2018) b. −log10(P-values) for enrichments in A calculated with hypergeometric tests. Horizontal line represents p-value of 0.05.

We next investigated the relationship between ΔPAU in humans and chimpanzees and the effect sizes for differences in protein translation between the species. (S. H. Wang et al., 2018). Considering the most differentially used 3’ UTR or intronic PAS per gene (**methods**), we identified a significant correlation between inter-species differences in translation and ΔPAU for 3’ UTR PAS, with a stronger correlation among genes with significant differences in both APA and translation (**Figure 5 - figure supplement 2**). As expected, and to some extent we view this as a control analysis, we did not identify a significant correlation between intronic PAS ΔPAU values and differences in translation (**Figure 5 - figure supplement 2**).

Given the apparent impact of PAS usage on protein translation, we next considered direct measurements of protein expression data from 3,391 genes in LCLs from humans and chimpanzees (Khan et al., 2013). Using summary statistics from this study, we found 1,263 genes to be differentially expressed at the protein level between the species (FDR of 5%). As the protein measurements are restricted to these 3,391 genes, we do not have enough power to ask if genes with inter-species differences in APA are also more likely to be differentially expressed at the protein level. However, we did find a positive correlation between the absolute value of 3’ UTR ΔPAU and the standardized number of ubiquitination sites for the same gene (Pearson’s correlation, R=0.15, *p*=5.0×10^−7^, **Figure 5 - figure supplement 3, methods**), consistent with the observation that 3’ UTR PAS are targets for the regulation of protein decay (Dubnikov et al., 2017; Ravid & Hochstrasser, 2008).Thus, we next focused on the 506 genes with significant inter-species differences in protein expression and an absence of corresponding differences in transcript expression levels that we also tested for differences in APA. Khan *et al*. reasonably hypothesized that inter-species differences in translation could account for the emergence of differences in protein expression levels when there are no regulatory differences at the RNA level, but they were unable to point to specific mechanisms. These genes are particularly interesting in the context of our current study, because APA which results in changes to 3’ UTR length may be more likely to result in differences in protein expression without affecting the expression level of the mRNA.

Indeed, we found 76 genes with inter-species differences in APA that are also differentially expressed at the protein but not at the RNA level between humans and chimpanzees (**Figure 6A,B**). In these 76 genes, inter-species differences in PAS usage are enriched at the 3’ UTR (**Figure 6 - figure supplement 1**). Finally, to assess whether APA contributes to differences in gene regulation by affecting translation efficiency or protein degradation, we asked whether genes with differential protein expression were also differentially translated. Of the 149 genes with significant differences in APA and protein expression, Wang *et al*. reported translation measurements for 142 (S. H. Wang et al., 2018). Only 34 genes displayed significant differences in translation efficiency, suggesting that isoform-specific post-translational modification of protein levels is largely responsible for protein-level differences (**Figure 6C,D**).

**Figure 6:**
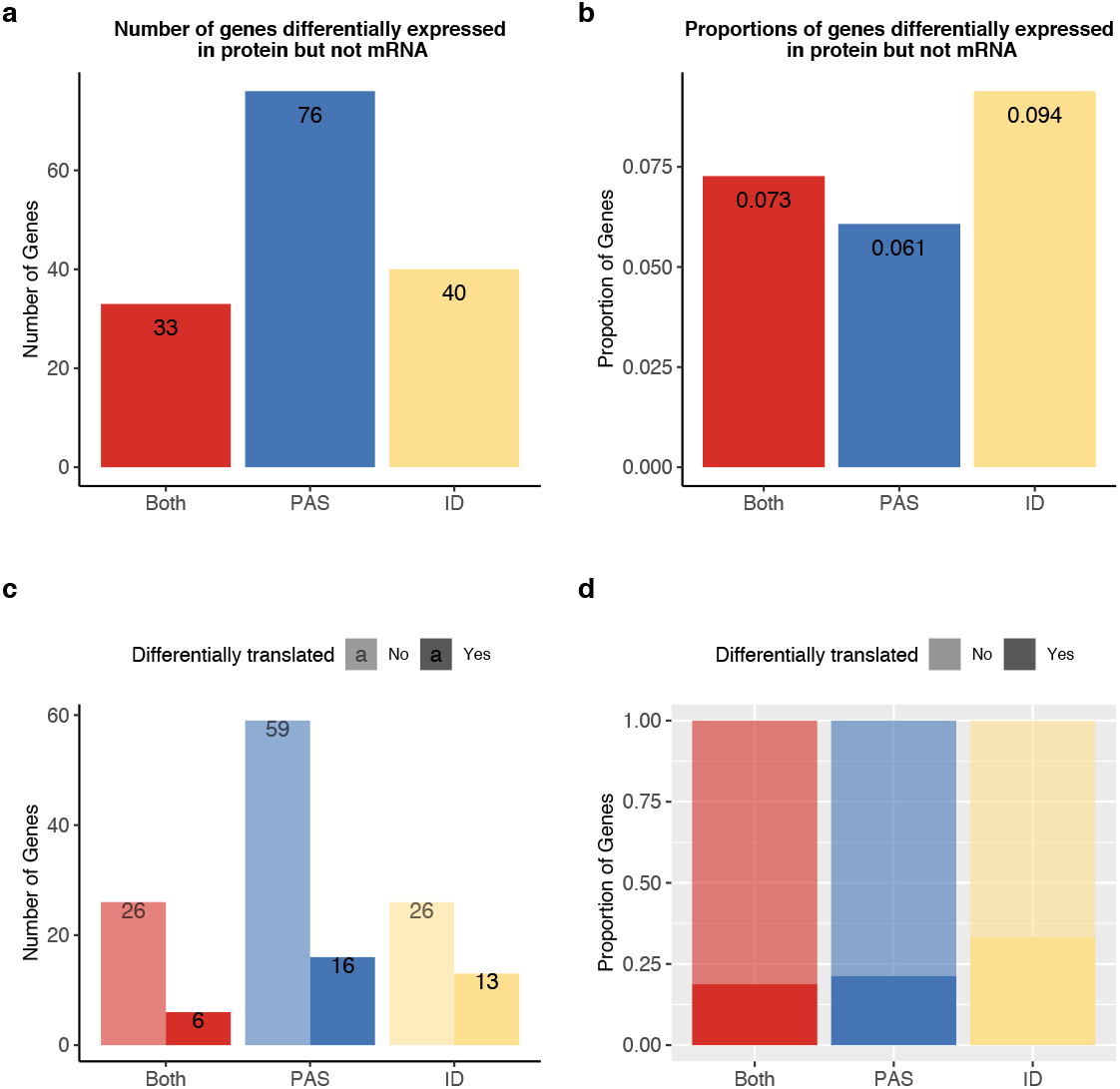
APA differences explain genes differentially expressed at protein level but not in mRNA. APA likely mediates functional differences post translationally. a. Number of genes with isoform level differences (ID), differences in APA at PAS level (PAS), or at both levels (Both) differentially expressed in protein (5% FDR), but not mRNA (5% FDR). Genes with differentially expressed protein reported in Khan et al. b. Proportion of genes with differential usage at PAS level (1,251 genes), isoform diversity level (426 genes) or both (454 genes), differentially expressed in protein (5% FDR), but not mRNA (5% FDR). c. Genes reported in a separated by genes differentially translated at 5% FDR. Differentially translated genes reported in Wang et al. (S. H. Wang et al., 2018) d. Genes differentially expressed in protein but not in mRNA, colored by differences in APA. Proportion of genes in the set differentially translated at 5% FDR.

## Discussion

Comparative primate functional genomic studies have contributed to our understanding of the gene regulatory processes that underlie genotype-phenotype relationships. A common goal of these studies is to understand the general properties and level of conservation of specific regulatory phenotypes, such as gene expression or DNA methylation levels. Multiple data types collected from the same cell lines or tissues can then be analyzed together to generate hypotheses about how gene regulatory processes contribute to inter-species differences in morphology, physiology, cognitive phenotypes, disease susceptibility, and other traits (Blake et al., 2020; Khan et al., 2013; Romero et al., 2018; S. H. Wang et al., 2018; Zhou et al., 2014). Moreover, unlike functional genomic studies within humans, comparative studies require only modest sample sizes to identify regulatory effects. This is because genetic variation between species is greater than genetic variation within species. Thus, regulatory differences that distinguish humans from other primates tend to have larger effect sizes than regulatory differences that distinguish between individual humans (Housman & Gilad 2020).

Importantly, inter-species differences identified using a comparative approach may be important not only for understanding primate evolution, but may implicate candidate loci for further investigation in humans. For example, genomic regions that are conserved in primates may point to loci that are likely to have negative functional consequences in humans, potentially with effects on disease risk (Housman et al., 2019). Identification of genomic regions under adaptation in humans is also critical, as they may point to causal mechanisms for human-specific traits, including diseases that are specific to humans (Gokhman et al., 2020; Ward & Gilad, 2019).

We characterized APA in human and chimpanzee LCLs to begin to understand the role that co-transcriptional mechanisms play in the evolution of gene regulation in primates. Our group has previously studied a variety of other gene regulatory phenotypes in primate LCLs (Cain et al., 2011; Khan et al., 2013; S. H. Wang et al., 2018; Zhou et al., 2014). We and others have demonstrated that gene regulatory phenotypes in these cell lines recapitulate many regulatory patterns seen in primary tissues (Çalışkan et al., 2011, 2014; Khaitovich et al., 2006). Not only did our use of primate LCLs allow us to circumvent many of the practical and ethical issues associated with primate research, but it also allowed us to integrate 3’ Seq and gene expression data from this study in the context of previously collected ribosome occupancy and protein expression data from primate LCLs (Khan et al., 2013; S. H. Wang et al., 2018). Together, these data allowed us to study the contribution of APA to inter-species differences in transcript and protein expression levels.

APA is an important molecular mechanism with regard to both the evolution of gene regulation and physiological traits. On a long-term evolutionary scale, both 3’ UTR length and the proportion of genes exhibiting APA have scaled with genome size and complexity. The expansion of APA is believed to have introduced biological complexity independent of an increase in the number of distinct genes (Mayr, 2016, 2017). As usage of multiple isoforms has been maintained, it is likely that distinct isoforms have divergent functions that are maintained by balancing selection. For example, APA facilitates post-transcriptional regulation of a *Drosophila Hox* gene through maintenance of two isoforms differentially targeted by multiple miRNAs (Patraquim et al., 2011). In turn, genome-wide changes in APA during differentiation of stem cells to terminal cell types directs isoform-specific gene regulation that is important for development in a range of species, including humans (Hilgers et al., 2011; Ji et al., 2009; Y. Li et al., 2012). Further, dysregulation of tumor suppressor genes through intronic polyadenylation is known to contribute to cancer pathogenesis (Dubbury et al., 2018; Lee et al., 2018). We hypothesize that a better understanding of APA in primates will aid in the understanding of APA evolution and its contribution to human-specific phenotypes.

### Alternative polyadenylation is mostly conserved, especially dominant sites

We measured APA from 3’ Seq data by calculating a ratio of isoforms terminating at one PAS compared to isoforms terminating at other PAS for the same gene. We then compared PAS usage ratios between species. To expand our understanding of APA conservation, we also calculated an isoform diversity statistic (**Simpson’s D**) for each gene in each species. Because Simpson’s D captures both the number of PAS isoforms and their usage, we were able to evaluate small regulatory changes spread across many PAS, rather than only focus on large changes at individual PAS. While previous studies have used Shannon’s Index to quantify isoform diversity (Pai et al., 2016; S. H. Wang et al., 2018), we found Simpson’s D to be less correlated with PAS number, making it less sensitive to the number of PAS per gene (**Figure 2 - figure supplement 5,6**). In addition, by placing more weight on dominant PAS, Simpson’s D more closely mirrors our current biological understanding of APA, wherein dominant PAS play a larger role in downstream gene regulation (Morris et al., 2014).

In general, we found that both individual PAS usage and isoform diversity are highly conserved between human and chimpanzee. Consistent with comparative studies of APA in humans and rodents, which used genomic synteny to identify conserved PAS, we found higher conservation among genes with a single PAS (Ara et al., 2006; R. Wang et al., 2018) and showed that sequence variation in PAS signal sites and the surrounding U-rich regions contributes to inter-species differences in APA (R. Wang et al., 2018). Because we characterized APA in closely related primates, our study provides additional insight into APA divergence at both the gene level and the species level, revealing functional changes that contribute to differences in downstream gene regulation. For example, we observed that when genes use one PAS markedly more often than others, said dominant PAS tends to be the same in both human and chimpanzee. It is likely that strong selection pressures have acted on these genes, resulting in continual usage of the same dominant isoform. This could imply that the dominant isoform is functionally important and alternative isoforms are potentially associated with reduced fitness. However, non-dominant isoforms also show evidence of conservation. Thus, it remains possible that there is a threshold at which the level of expression of the alternative isoforms begins to impede gene function.

### Alternative polyadenylation is associated with gene expression divergence

Our study also revealed that the majority of differentially used PAS between species are located in 3’ UTRs. We showed that, across species, increased intronic PAS usage is associated with increased mRNA expression levels, while increased 3’ UTR PAS usage is correlated with a decrease in mRNA expression. In a previous study, we found that human apaQTL alleles associated with increased intronic PAS usage were correlated with *decreased* mRNA expression levels (Mittleman et al., 2020). This is not the first molecular phenotype wherein a within-species study revealed alternative regulatory models compared to an inter-species analysis. For example, Pai *et al*. reported tissue-specific differential methylation to be almost exclusively inversely correlated with gene expression patterns between human and chimpanzee (Enard et al., 2004; Pai et al., 2011; Weber et al., 2007), whereas Banovich *et al.* discovered genetic variation associated with DNA methylation variation that was both directly and inversely correlated with eQTLs. At this time, we can’t provide evidence for a mechanistic explanation for these contradictory observations. We hypothesize the following: Transcripts terminating in introns are likely subject to nonstop decay (**NSD**). By studying APA variation within humans we probably captured the effects of intronic termination. Across species, however, increased intronic PAS may simply track the overall expression level of the gene. Specifically, increased usage of intronic PAS may result in truncated isoforms that do not contain 3’ UTR *cis* regulatory elements that would normally signal mRNA decay. If these truncated isoforms are no longer targets of mRNA decay, this could cause some of these genes to appear up-regulated. We hypothesize that the within-species effects related to NSD were likely overshadowed by inter-species differences, which typically have much larger effect sizes than differences observed within a population (Housman & Gilad, 2020).

Alternatively, the large effect sizes for differential usage of 3’ UTR PAS could also be driving the relationship between differential APA and differential expression. In line with this hypothesis, we found increases in AU destabilizing elements and ubiquitination marks for genes with divergent 3’ UTR PAS. However, since PAS usage is calculated as a ratio, we may have detected changes in intronic PAS usage solely as a mathematical consequence of changes in 3’ UTR PAS. Functional follow-up on the genes with PAS detected as differentially used between and within species would be necessary to explore the relative importance of each of these regulatory pathways and to disentangle the results from both studies.

In past studies, we and others have estimated the proportion of variation in gene expression explained by different regulatory mechanism. For example, we have previously tested for differences in expression before and after accounting for another regulatory mechanism or used formal mediation analyses (Blake et al., 2020; Blekhman et al., 2009; Cain et al., 2011; Eres et al., 2019). We would have liked to perform similar analysis in the current study, regarding the role that APA plays in overall regulatory divergence between the species. However, APA was measured using ratios of alternative mRNA isoforms and thus, effect sizes for APA and differential expression are on different scales and we cannot use a standard mediation approach to formally calculate the proportion of expression variation explained by APA. That said, we are generally convinced that APA contributes to differences in mRNA expression overall because 764 of 3,796 (20.1%) differentially expressed genes also have significant differences in APA.

### Inter-species differences in APA explain protein-specific regulatory divergence

Though genes are ultimately expressed as proteins, many studies (including current studies by our group) still measure mRNA expression as an implicit proxy for protein expression levels. As a justification for this approach, we typically point to the fact that after accounting for technical considerations, the correlation between mRNA levels and protein abundance is quite high genome-wide, specifically, across genes (Buccitelli & Selbach, 2020; Csárdi et al., 2015). However, we also know that at the level of a single gene, across individuals or tissues, the correlation between mRNA and protein measurements tends to be much lower (Battle et al., 2015; Buccitelli & Selbach, 2020). This suggests that a number of molecular mechanisms decouple mRNA and protein expression levels post-transcriptionally. Clearly, we do not yet fully understand the post-transcriptional and translational mechanisms that shape the proteome (Buccitelli & Selbach, 2020).

Within human populations and between primates, there is large number of genes that are differentially expressed at the mRNA level, but not as proteins. By directly measuring translation levels for these genes, previous studies have proposed that post-translational protein buffering can explain the decreased variation at the protein level (Battle et al., 2015; S. H. Wang et al., 2018). Conversely, there are also genes that are more variable at the protein level than at the mRNA levels (Battle et al., 2015; Chick et al., 2016; Khan et al., 2013). Our previous work demonstrated that some protein-specific QTLs are also highly correlated with differences in APA (Mittleman et al., 2020). Considered all of these observations, we expanded this analysis and demonstrated that genes that are differentially expressed as proteins, but not at the mRNA level, between human and chimpanzee, tend to have divergent APA patterns. We also found that the divergent protein levels are likely due to post-translational molecular mechanisms. While we cannot directly test the mechanism here, we hypothesize that APA could lead to variation in protein levels as a consequence of protein autoregulation, by differentially including RNA and protein binding motifs (Buccitelli & Selbach, 2020; de Bie & Ciechanover, 2011; Müller-McNicoll et al., 2019). Alternatively, APA could contribute to temporal and spatial differences in protein expression, which would affect our ability to quantify protein with traditional techniques (Buccitelli & Selbach, 2020; Tian & Manley, 2017).

In conclusion, a better understanding of co-transcriptional gene regulatory mechanisms, such as APA, may point to additional mechanisms contributing to the decoupling of mRNA and protein abundance and more generally, enhance our understanding of how variation percolates through genetic variants, mRNA, and protein to ultimately affect human phenotypic diversity.

## Methods

### Cell culture and collections

We grew 6 human and 6 chimpanzee Epstein-Bar virus transformed lymphoblastoid cell lines (LCLs) in glutamine depleted RPMI [RPMI 1640 1X from Corning (15-040-CM)], completed with 20% FBS, 2mM GlutaMax [Gibco (35050-061)], 100 IU/ml Penicillin, and 100 ug/mL Streptomycin. We cultured all cells at 37°C at 5% CO2. We passaged each cell line a minimum of 3 times then maintained cells at 1 × 10^6^ cell per mL in preparation for collection. Cell line numbers and details can be found in **Supplementary table 2**. The human lines were derived from Yoruba individuals collected as part of the HapMap project and can be ordered through the Coriell Institute (International HapMap Consortium, 2005). Chimpanzee LCLs were originally transformed from individuals from the New Iberia Research Center (University of Louisiana at Lafayette), Coriell IPBR repository and Arizona State University (Khan et al., 2013). The cell lines have previously been used for similar studies of primate gene regulation (Cain et al., 2011; Khan et al., 2013; S. H. Wang et al., 2018; Zhou et al., 2014).

Once all cells lines reached 1 × 10^6^ cell per mL, we used the collection and RNA extraction method detailed in Mittleman et al. to extract whole cell and nuclear mRNA. Briefly, we collected 30 million cells in two 15 million cell aliquots. We extracted nuclei from one aliquot per line using the nuclear isolation protocol outlined by Mayer and Churchman (Mayer & Churchman, 2016). We extracted mRNA in two fraction- and species-matched batches, using the miRNeasy kit (Qiagen) according to manufacture instructions, including the DNase step to remove genomic DNA. We quantified mRNA and tested quality using a nanodrop. Details of mRNA processing for each line, including concentrations and quality can be found in **Supplementary table 2**.

### 3’ Sequencing to identify PAS and quantify site usage

We generated 3’ sequencing (**3’ Seq**) libraries from whole cell and nuclear-isolated mRNA from 6 chimpanzee and 6 human individuals using the QuantSeq Rev 3’ mRNA-Seq Library Prep Kit (Moll et al., 2014) according to the manufacturer’s instructions. We sequenced all libraries on the Illumina NextSeq500 at the University of Chicago Genomics Core facility using single-end 50 bp sequencing.

We mapped human 3’ Seq libraries to GRCh38 (Schneider et al., 2017) and chimpanzee libraries to panTro6 (Chimpanzee Sequencing and Analysis Consortium, 2005) using the STAR RNA-seq aligner with default settings (Dobin et al., 2013). Similar to our previous work, we removed reads with evidence of internal priming resulting from the poly(dT) primer. We filtered reads proceeded by 6 As or 7 of 10 A’s in the base pairs directly upstream of the mapped location (Mittleman et al., 2020; Sheppard et al., 2013; Tian et al., 2005). To ensure that differences in low quality bases would not bias our results, we treated any N in the genome annotation as an A. All raw read counts, mapped read counts, and filtered read counts can be found in **Supplemental table 2**.

We first identified an inclusive set of PAS in each species separately. We used the same in-house peak caller described in Mittleman et al (Mittleman et al., 2020), annotating each PAS as the most 3’ base in each peak. The initial PAS set included 340,023 in human and 303,249 in chimps. We extended PAS 100 bp upstream and 100 bp downstream and used a reciprocal liftover pipeline to identify an inclusive set of orthologous PAS. We downloaded chain files from UCSC genome browser (W. James Kent et al., 2002). Details of the pipeline and number of PAS passing each step can be found in **Figure 1 - figure supplement 11**.

Due to gene annotation differences between species, we annotated all orthologous PAS to the human NCBI RefSeq annotation downloaded from UCSC genome browser (Pruitt et al., 2005). We used a hierarchical model to assign PAS to genic locations (Lin et al., 2012; Mittleman et al., 2020). We prioritized annotations in the following order: 3’ UTRs (UTR5), 5kb downstream of genes (end), exons (cds), 5’ UTRs (UTR5), and introns (intron). We quantified reads mapping to each annotated PAS for each individual in both the total RNA libraries and nuclear RNA libraries using featureCounts with the -s strand specificity flag (Liao et al., 2014). We calculated usage for each PAS in each library as a ratio of reads mapping to the PAS divided by the number of reads mapping to any PAS in the same gene (**Figure 1 - figure supplement 2)**. We implemented two filtering steps to remove PAS with ratios likely biased by low site count or low gene count separately in each fraction.

Next, we filtered out sites with less than 5% usage in both species in the nuclear fraction. We then merged nuclear counts across all PAS in each gene. We removed PAS in genes not passing a cutoff of log2(CPM)>2 in at least 8 of the 12 individuals. After applying these filters, we were left with 44,432 PAS. As a quality control metric, we compared PAS usage calculated from the nuclear fraction to PAS usage calculated from whole cell fraction for each individual (we used the same methods to identify and quantify PAS usage in the whole cell 3’ Seq data). We expected a high correlation between PAS usage in each fraction. Further, we expected a similar correlation in human and chimpanzee individuals (Mittleman et al., 2020). Human individual NA18499 had significantly lower across-species correlation than the other individuals and was therefore removed from the analysis (**Figure 1 - figure supplement 3**).

To ensure gene expression level did not introduce ascertainment bias, we tested the relationship between PAS number and normalized gene expression. In both species, number of PAS is negatively correlated with normalized gene expression (Human: Pearson correlation =-0.19, p< 2.2×10^−16^, Chimp: Pearson correlation=-0.17, p< 2.2×10^−16^, **Figure 1 - figure supplement 6**). We expected species to contribute the most amount of variation to PAS usage. We ran PCA on the filtered nuclear PAS usage. PC1 accounts for 41.8% of the variation and is highly correlated with species (R^2^ =0.68). PC2 accounts for 13.1% of the variation and is moderately correlated with RNA extraction technician (R^2^=0.38) and extraction day (R^2^=0.28). As both of these variables are balanced with respect to species, we do not believe they bias the results (**Figure 1 - figure supplement 5**). We identified 302 sites used at a rate of 5% or greater in humans and 0% in chimpanzees, which we designated as human-specific. We identified 357 sites used at a rate of 5% or greater in chimpanzee and 0% in humans, which we designated as chimp-specific.

We acknowledge the possibility that unlifted PAS may affect downstream analyses; therefore, we removed genes for which PAS ratios may be affected. Specifically, we annotated and calculated usage for the human PAS, including the 10,077 PAS that do not reciprocally lift to the chimpanzee genome. After removing PAS in genes previously identified as lowly expressed and PAS with usage below 5%, 386 PAS in 353 genes remain (**Figure 2 - figure supplement 8**). We removed these 353 genes and recreated main figures 3–6 (**Figure 3 - figure supplement 3, Figure 4 - figure supplement 1, Figure 5 – Figure Supplement 4, Figure 6 - figure supplement 2**).

### Orthologous 3’ UTRs

We identified a set of orthologous UTRs using the orthologous exon file described the Differential Expression section of the methods. We merged all regions annotated as 3’ UTR by gene. If a gene had multiple non-continuous annotations, we selected the most 3’ region as the orthologous UTR. We used deepTools compute matrix and plotHeatmap functions to plot merged human and chimpanzee reads along the orthologous 3’ UTR set (**Figure 1 - figure supplement 1**) (Ramírez et al., 2016) For all genes with PAS only in 3’ UTRs, we assigned PAS to single, first, middle, and last, as previously described (R. Wang et al., 2018).

### Analysis of sequence conservation around PAS

We used phyloP scores to measure sequence level conservation. We downloaded the hg38 100-way vertebrate PhyloP bigwig file from the UCSC table browser (Pollard et al., 2010). We computed scores for PAS regions as well as 200 bp intervals by taking the mean of the base pair scores. We removed any region with missing data from the analysis. We tested for differences in mean phloP scores using Wilcoxon rank sum tests.

We tested for presence of the polyadenylation signal site motif in the 200 bp PAS regions. We used the bedtools nuc tool with the strand-specific flag to test for presence of each of the 12 previously annotated motifs for each PAS in both species (Beaudoing et al., 2000; Quinlan & Hall, 2010). If a PAS had multiple motifs, we used a hierarchical model to choose the site based on the number of PAS with each identified motif (order: AATAAA, ATTAAA, AAAAAG, AAAAAA, TATAAA, AATATA, AGTAAA, AATACA, GATAAA, AATAGA, CATAAA, ACTAAA). The proportion of PAS with each signal site motif matched across species (**Figure 2**). To ask if presence or absence of a signal site explained species specificity or site-level differences, we restricted our analysis to the top two signal sites. These two motifs are the only sites where presence of a signal is associated with increased usage of the site in both species (**Figure 1 - figure supplement 9**). For the 359 PAS with one of these two signal sites present only in chimpanzees, average usage was higher in chimpanzees than in humans (p=0.025). For the 361 PAS with one of these two signal sites present only in humans, average usage was higher in humans (p=2.0×10^−4^). We used hypergeometric tests to evaluate enrichment of differentially used PAS and species-specific PAS in the set of PAS with signal sites in only one species.

We also examined the proportion of U nucleotides in each PAS region. We used the bedtools nuc with the -s flag for strand specificity (Quinlan & Hall, 2010). We tested if PAS with differences in U content are enriched for differentially used PAS using a hypergeometric test.

### Differential APA

#### PAS level differences

We quantified reads mapping to each PAS using the featureCounts tool with the -s strand specificity tool (Liao et al., 2014). We tested for site-level differences between human and chimpanzee using the leafcutter leafcutter_ds.R tool with standard settings (Li 2018). We tested for differences in both the total and nuclear fractions. We tested 43,038 PAS in 8,422 genes in the nuclear fraction and 41,914 PAS in 8,333 genes in the total fraction. We classified PAS as differentially used if the gene reached significance at 5% FDR and the PAS had a ΔPAU greater than 20% (absolute value (ΔPAU) > 0.2). A negative ΔPAU indicates increased usage in chimpanzees and ΔPAU indicates increased usage in humans. The top PAS per gene is the PAS with the most significant difference between species; ties were broken using mean usage for all individuals in both species.

#### Isoform diversity differences

We calculated Shannon Information content 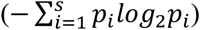 and Simpson’s Index 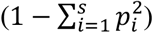 using mean usage of each PAS in humans and chimpanzees, where 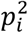 is the usage of the i^th^ of *s* sites in the gene. We used Simpson’s Index to assess isoform diversity because the correlation between Simpson’s Index and number of PAS is lower than the correlation between Shannon Information content and the number of PAS per gene (**Figure 2 - figure supplement 5,6)**. To identify genes with differences in isoform diversity, we recalculated Simpson’s index per gene per individual and tested for differences between species with Wilcoxon tests. We reported genes with differences at 5% FDR.

#### Conservation of dominant PAS

We consider a gene to have a dominant PAS if the within species average usage of the top used PAS is greater than the second most used site by 0.4. We reported results for cutoffs between 0.1 and 0.9. If a gene had a dominant PAS in either species, we included the top used site for both species when testing if genes use the same or different dominant PAS between species. We tested for enrichment of genes using the same or different dominant PAS with differentially expressed genes using hypergeometric tests.

### Differential expression analysis

We generated unstranded RNA-seq libraries using the Illumina TruSeq Total RNA kit according to the manufacturer’s instructions using the total mRNA collected from all 12 individuals (Illumina, San Diego, CA, USA). We sequenced RNA-seq libraries at the University of Chicago Genomics Core facility using the single-end 50 bp protocol on one lane of the Illumina HiSeq 4000 machine. RNA quality and concentration at the time of library prep and number of sequenced reads per library are available in **Supplementary table 3**. We mapped the human libraries to GRhg38 (Schneider et al., 2017) and chimpanzee libraries to panTro6 and quantified reads mapping to orthologous exons.

To generate an updated orthologous exon file for the most recent chimpanzee genome assembly (panTro6) we followed the procedure reported in Pavlovic et al. with slight modifications (Pavlovic et al., 2018). We started with human (GRCh38) exon definitions from Ensembl version 98. We filtered this set of definitions for biotypes ‘protein_coding’ using the command mkgtf from cellranger (10XGenomics). We then removed exon segments that were in exon definitions for multiple genes. This broke some exons into smaller unique exons. We then removed exons smaller than 10 bp. We took the final set of exons (1,371,917 exons from 20,338 genes) and extracted their sequences from the genome Ensembl GRCh38.p12. We used BLAT V. 35 to identify orthologous sequences within the chimpanzee genome (panTro6) (W. J. Kent, 2002). We removed hits with indels larger than 25 bp (using a function blatOutIndelIdent from https://bitbucket.org/ee_reh_neh/orthoexon). We then extracted the panTro6 sequences that had the highest sequence identity. We ran BLAT on this orthologous exon set to find matches in both the human and chimpanzee genomes. We removed exons that did not return the original location in humans or chimpanzees, as well as exons that mapped to multiple places with higher than 90% sequence identity. We removed exons from different human genes that mapped to overlapping regions in the chimpanzee genome. Finally, we removed exons that mapped to a different contig than the majority of exons from each gene. This resulted in a set of 1,250,820 orthologous exons from 19,515 genes.

We mapped on average 18.6 million reads to orthologous exons. We collapsed orthologous exons to quantify raw gene expression for each gene in each individual. We standardized counts and filtered out genes that did not pass the criteris of log2(CPM) values greater than 1 in 8 of the 12 individuals. To prepare counts for differential expression modeling we used the Voom function with the quantile normalization method in the limma R package (Ritchie et al., 2015). We used PCA to test for batch effects. PC1 explains 35.1% of the variation and is highly correlated with species (R^2^= 0.98) (**Figure 3 - figure supplement 4)**. Collected metadata such as the percent of live cells at collection, cell concentration at collection, RIN score, and RNA concentration do not segregate by species (**Figure 3 - figure supplement 4**). We modeled species as a fixed effect and called genes as differentially expressed at a 5% FDR. The results from our differential expression analysis, including effect sizes and significance values are available in **Supplementary Table 3.**

### Integration of translation and protein data

We downloaded differentially translated genes and their effect sizes from Additional file 5 of Wang et al 2018 (S. H. Wang et al., 2018). Wang et al. modeled differential translation using ribosome profiling of 4 human, 4 chimpanzee, and 4 rhesus macaque LCLs. For all integrations, we conditioned on the 6,407 genes tested in the Wang et al. study and in our APA analysis. We tested for enrichments using a one-sided hypergeometric test implemented in R. We tested for correlations in effect sizes between site level ΔPAU and translation HvC effect sizes by first filtering for the top PAS (see top PAS method above, **Figure 5 - figure supplement 2**). We report Pearson’s correlations calculated in R.

We downloaded differential protein level genes, effect sizes, and directional selection classifications from **table S4** of Khan et al. 2013 (Khan et al., 2013). Khan et al. modeled differential protein expression of 3,390 genes using high resolution mass spectrometry of stable isotope labeling by amino acids in cell culture (SILAC) collected from 5 human, 5 chimpanzee, and 5 rhesus macaque LCLs.

### Supplemental functional data

We downloaded human protein length (in number of amino acids) for proteins annotated as reviewed for high confidence from UniProtKB (The UniProt Consortium, 2019). We downloaded ubiquitination protein modification data from PhoshoSitePlus version 050320 (Hornbeck et al., 2015). For all analyses in which we used interaction or ubiquitination data, we normalized the values by number of amino acids. To identify 3’ UTR AU-rich elements in human RefSeq annotated 3’ UTRs, we used the transcriptome_properties.py script published in Floor and Doudna 2016, available at https://github.com/stephenfloor/tripseq-analysis, with the ‒au-elements flag (Floor & Doudna, 2016). According to Floor and Doudna 2016, the fraction of AU-elements is the percentage of the 3’ UTR with repeating AU elements of 5nt or more (Floor & Doudna, 2016).

## Acknowledgements

We thank N. Gonzales for comments on the manuscript. We thank Y. Li, M. Ward, and G. Housman for useful discussion. Funding: This work was supported by the US National Institutes of Health (R01HG010772 and R35GM13172 to Y.G). B.E.M. supported by T32 GM09197 to the University of Chicago and F31HL149259 to B.E.M. from National Heart, Lung, And Blood Institute of the National Institutes of Health. SP was in part supported by the National Center for Advancing Translational Sciences of the NIH (K12 HL119995). This work was completed in part with resources provided by the University of Chicago Research Computing Center.

## Author Contributions

B.E.M. conceived the project with help from Y.G and S.P. B.E.M., S.W., C.C. and S.P. performed the experiments. B.E.M performed the analysis. K.B. curated orthologous exon file. B.E.M. drafted the manuscript with input from Y.G. and S.P. S.P. and Y.G supervised the project.

## Data and material availability

All scripts and analysis pipelines can be found at https://brimittleman.github.io/Comparative_APA/index.html. FastQ files and PAS annotations are available at GEO under accession **GSE155245**.

## Supplemental Figures

**Figure 1 - figure supplement 1:**
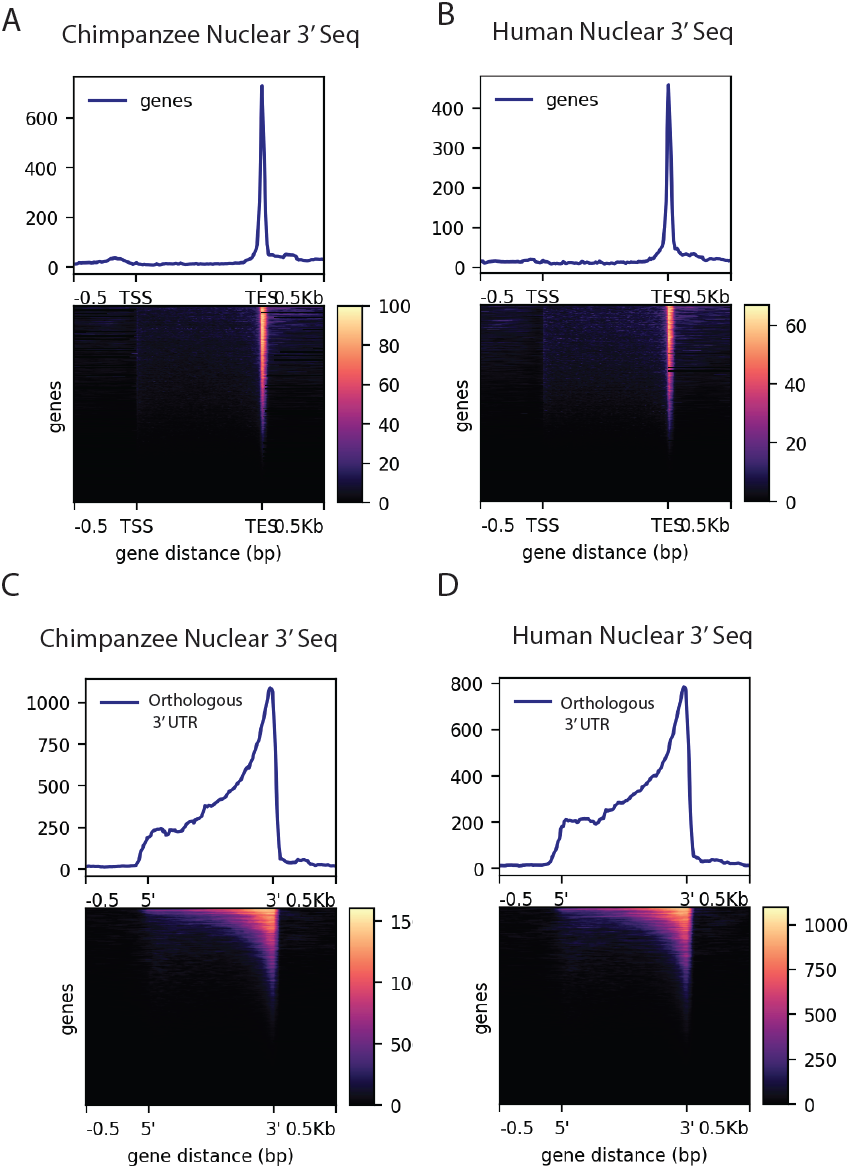
Density of merged human and chimpanzee 3’ Seq. A. Coverage of 6 chimpanzee, nuclear 3’ seq reads along Refseq transcripts B. Coverage of 5 human, nuclear 3’seq reads along Refseq transcripts C. Coverage of 6 chimpanzee, nuclear 3’ seq reads orthologous 3’ UTRs D. Coverage of 5 human, nuclear 3’ seq reads orthologous 3’ UTRs}

**Figure 1 - figure supplement 2:**
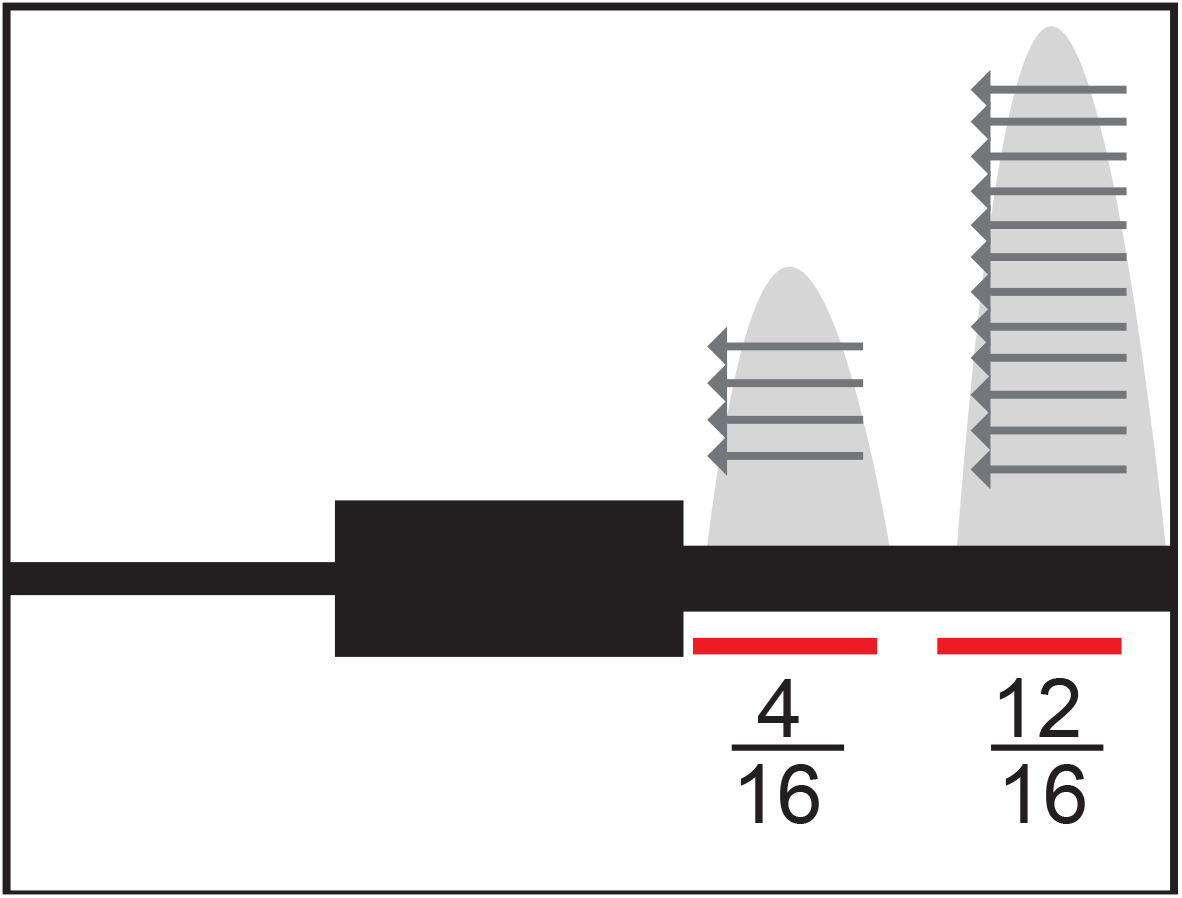
Model representation of usage calculation Representation of PAS usage calculation. Usage is a ratio of reads at each PAS to the number of reads mapping to any PAS in the same gene. Adapted from Mittleman *et al.*

**Figure 1 - figure supplement 3:**
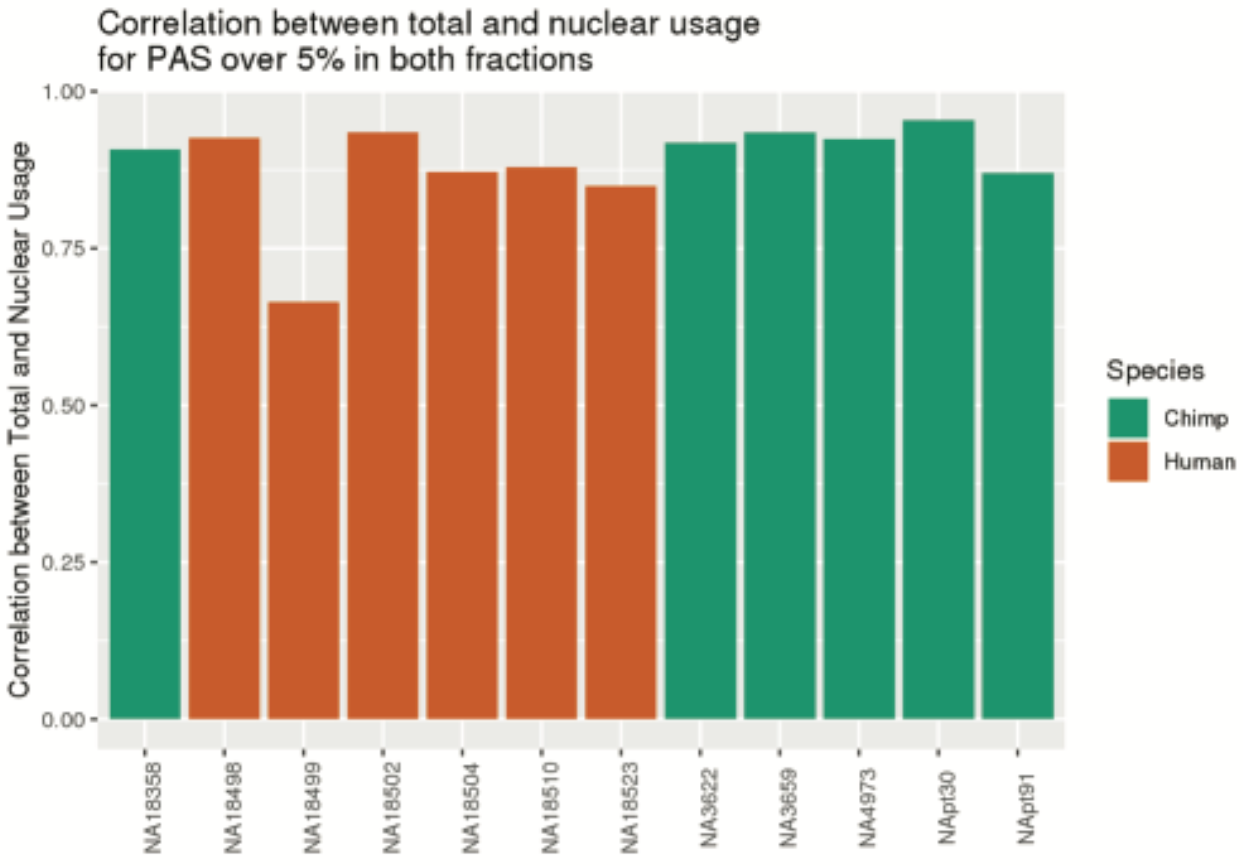
NA18499 removed from analysis due to low correlation between fractions. Pearson’s correlation between PAS usage calculated using nuclear 3’ seq libraries and total mRNA 3’ seq libraries, calculated using sites reaching 5% in one species in both fractions. (Plot from previous git commit 30ff122 on Aril 9, 2020, https://github.com/brimittleman/Comparative_APA)

**Figure 1 - figure supplement 4:**
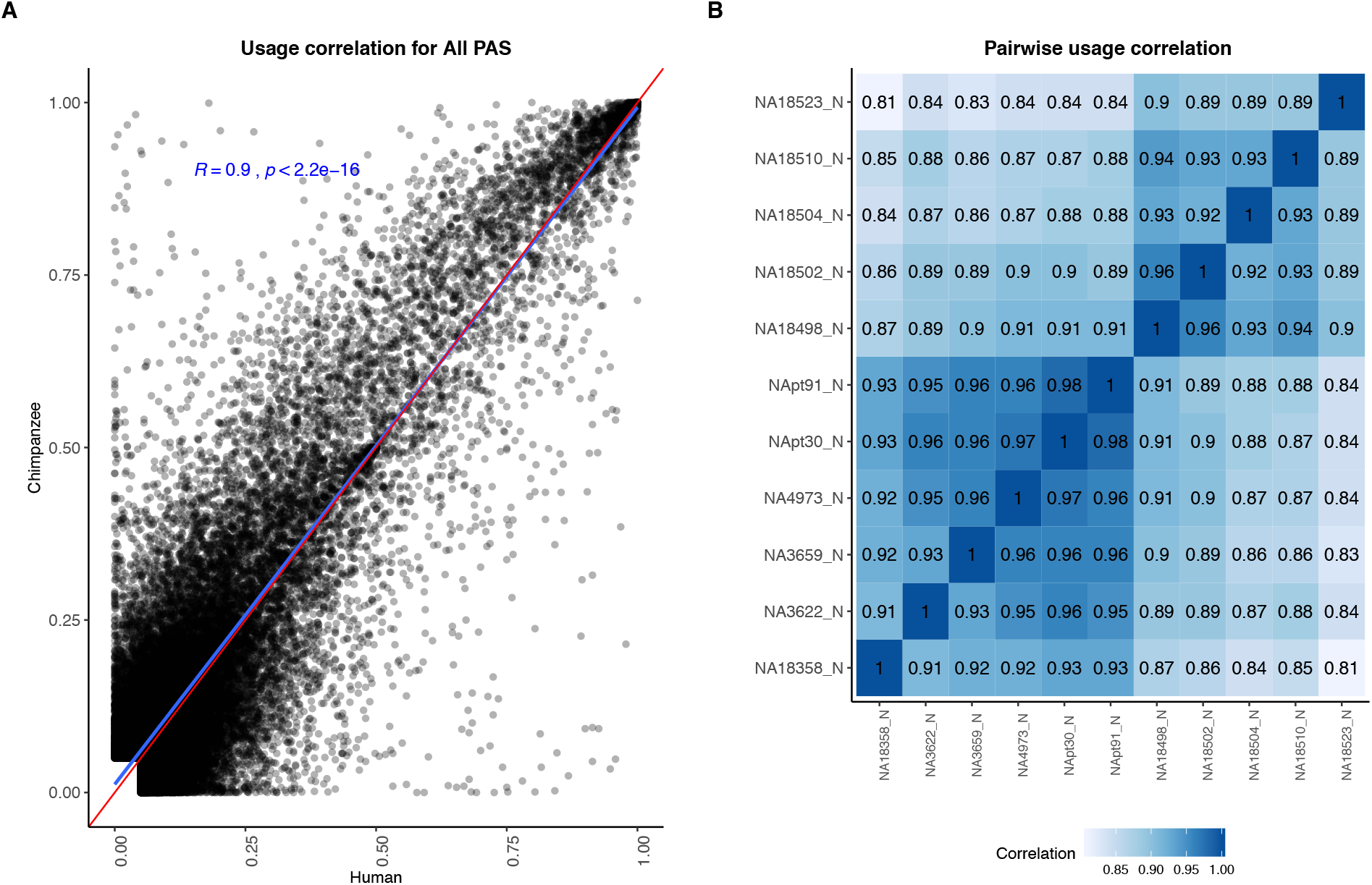
PAS usage is highly correlated across species. A. Correlation between human and chimpanzee PAS usage for 44,432 PAS. Red line is a 1:1 line. Linear regression line and Pearson’s correlation plotted in blue. B. Pairwise correlation for human and chimpanzee PAS usage.

**Figure 1 - figure supplement 5:**
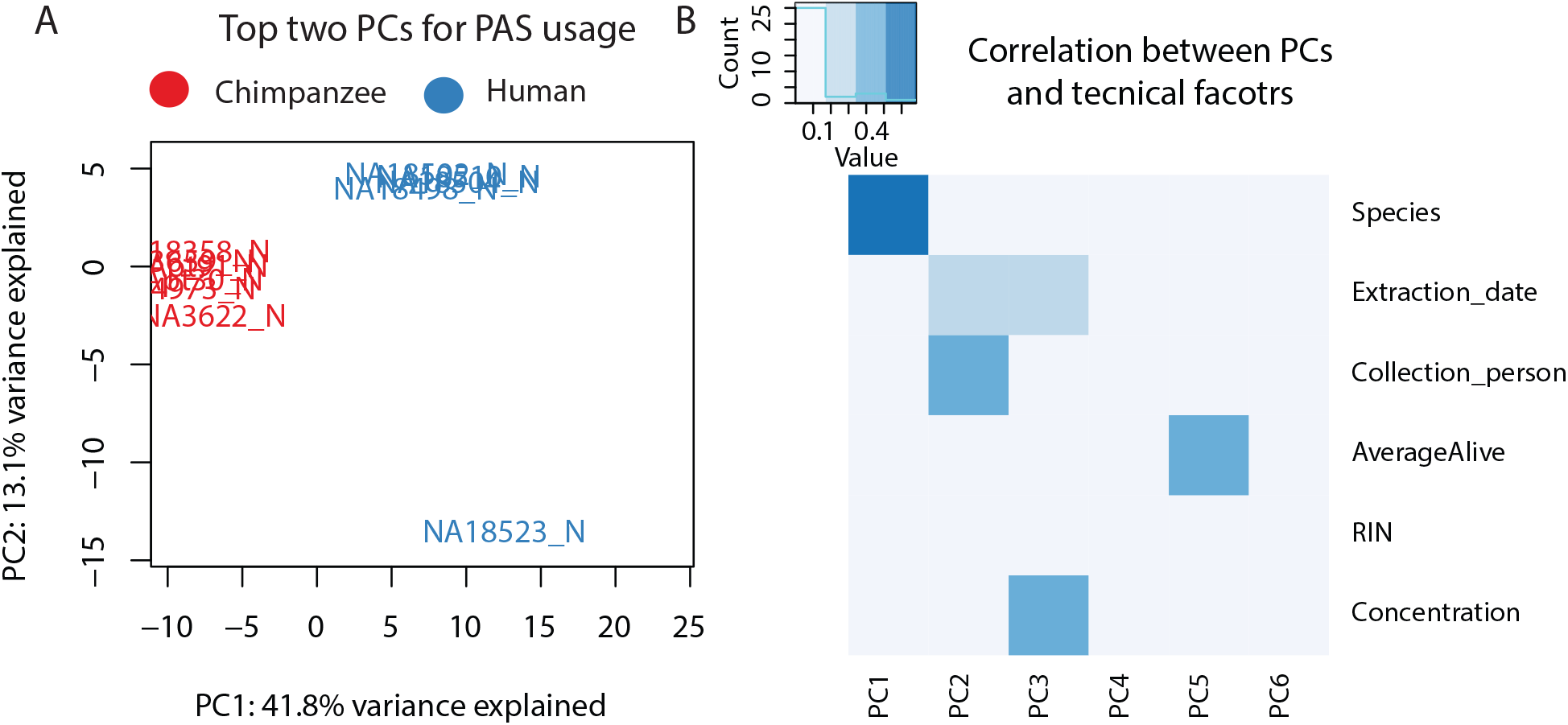
Variation in PAS usage. A. Plot of first two principal components (PCs) calculated by a principal component analysis on PAS usage (44,432 PAS). Chimpanzee samples are shown in red and human samples are shown in blue B. Heatmap representing correlation between technical factors and PCs. Y axis factors include: Species, Extraction date, Collection person, AverageAlive (average of two live dead calculations at time of collection), RIN score, RNA concentration. Explanation of factors and values in Supplemental Table 1

**Figure 1 - figure supplement 6:**
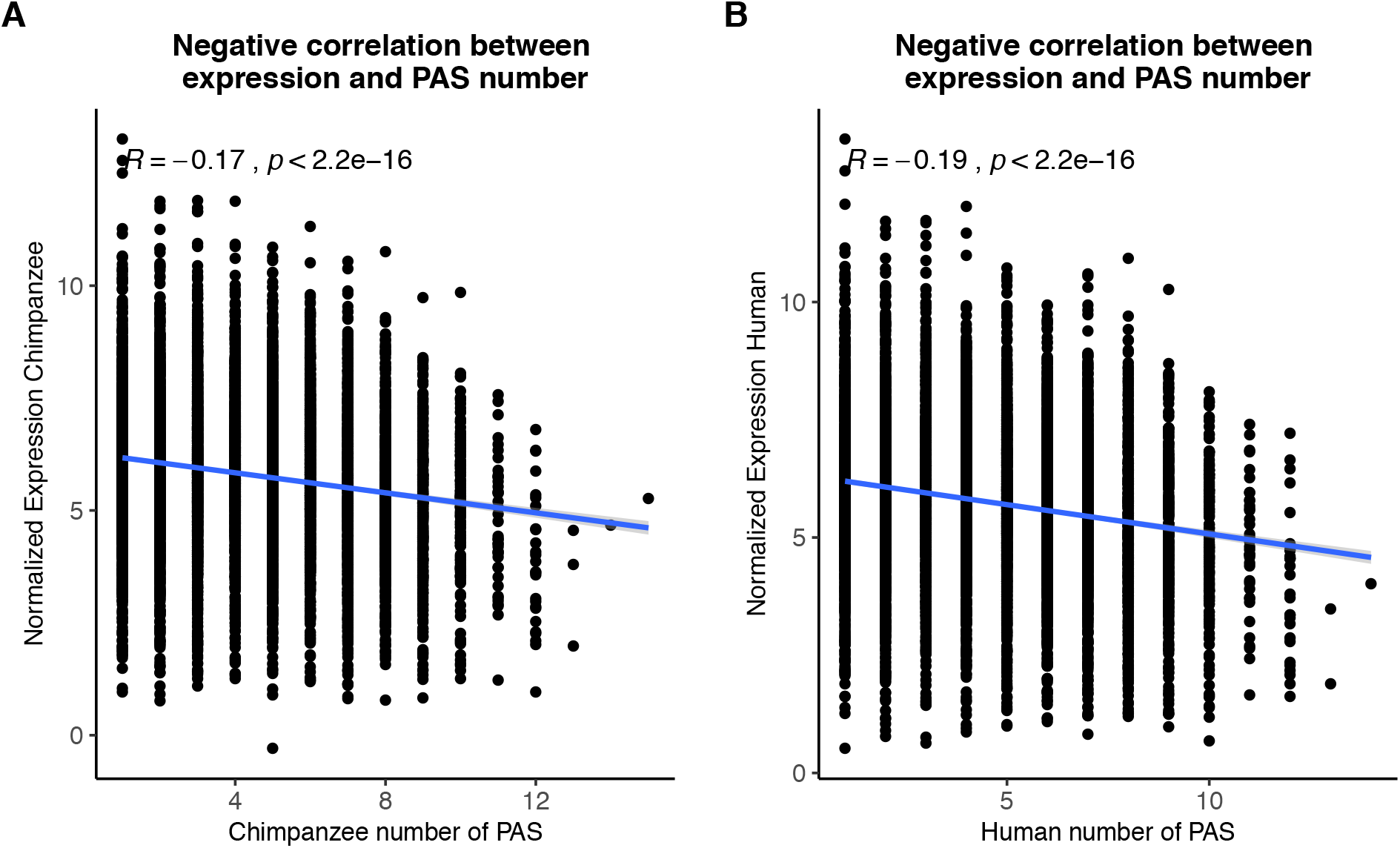
PAS detection likely not biased by expression level. A. Normalized gene expression plotted against the number of PAS detected at 5% usage in chimpanzee. B. Normalized gene expression plotted against the number of PAS detected at 5% usage in human. The R package ggpubr was used to plot linear regression lines and calculate Pearson’s correlations.

**Figure 1 - figure supplement 7:**
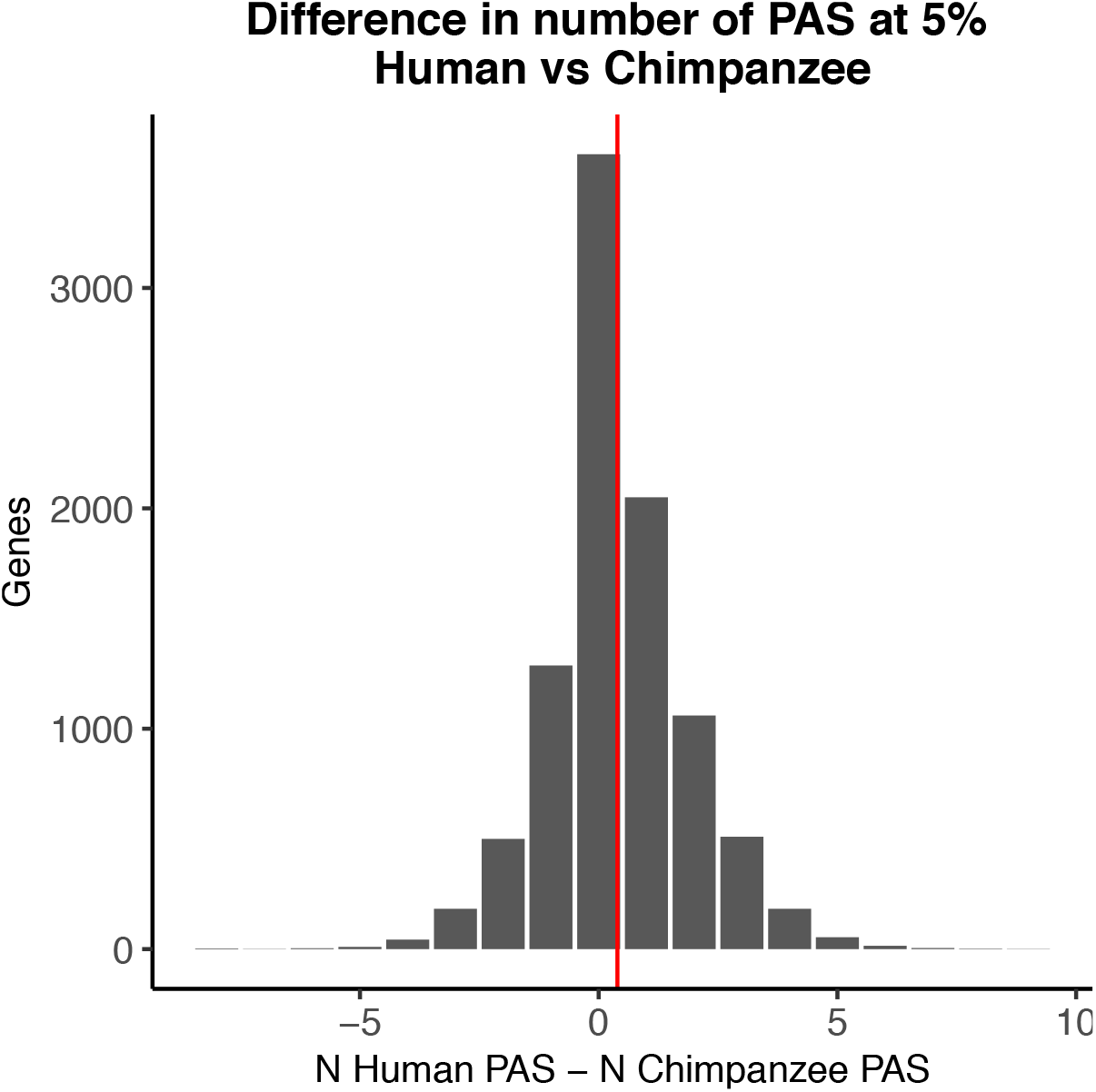
PAS detection likely not biased by species. Histogram of the number of PAS detected at 5% usage in human minus the number of 5% usage in chimpanzees. Red vertical line represents mean difference (0.39)

**Figure 1 - figure supplement 8:**
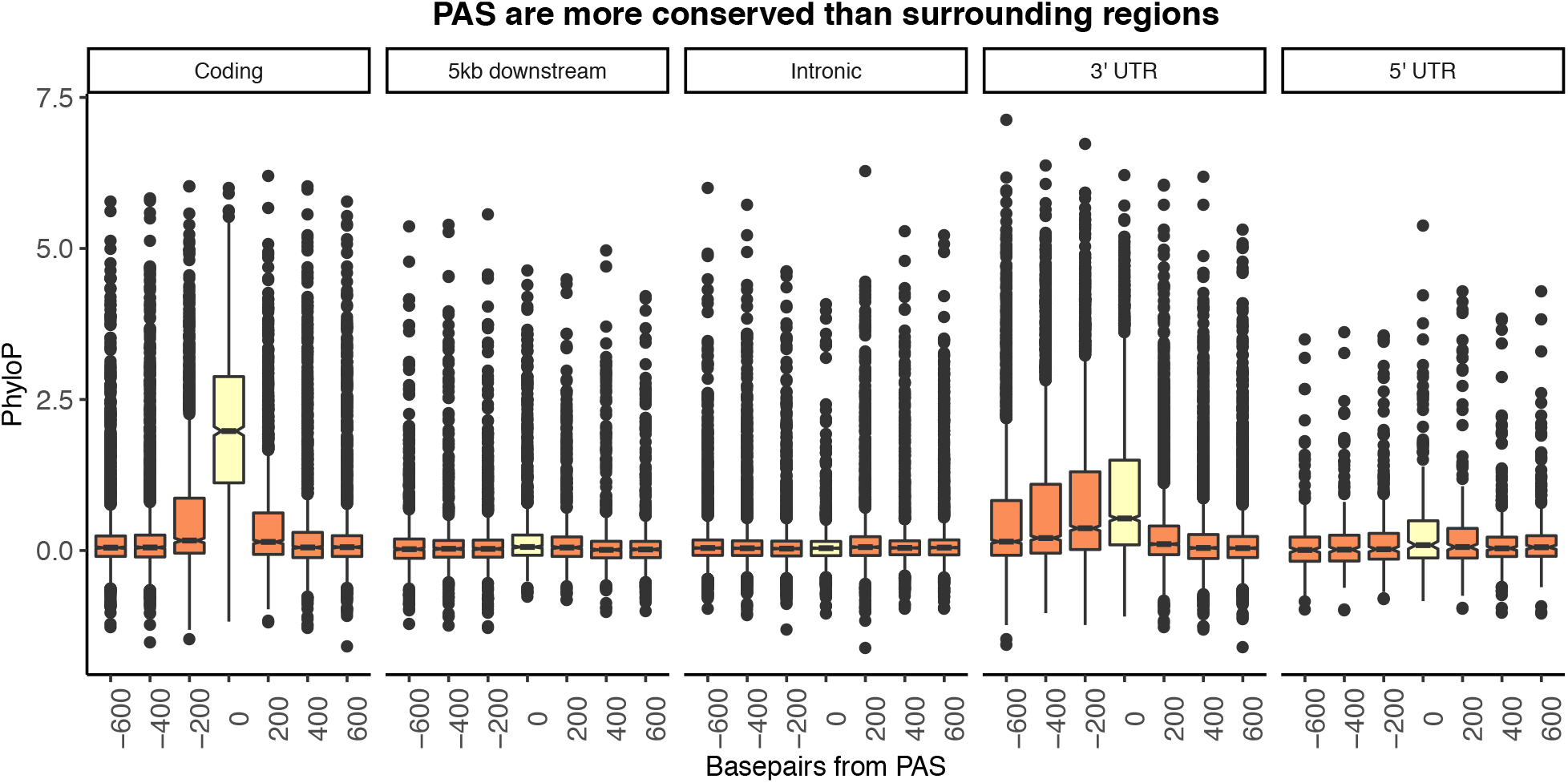
Figure 1B separated by genic location. Mean PhyloP scores for PAS regions (yellow) and 200 base pair bins upstream and downstream of PAS (orange). A one-sided Wilcoxon test was used to test for increased PhyloP in PAS regions (Coding region: p < 2.2×10^-16^, 5 kb downstream of genes: p = 7.02×10^-6^, intron: p=0.99, 3’ UTR: p < 2.2×10^−16^, 5’ UTR: p = 0.011).

**Figure 1 - figure supplement 9:**
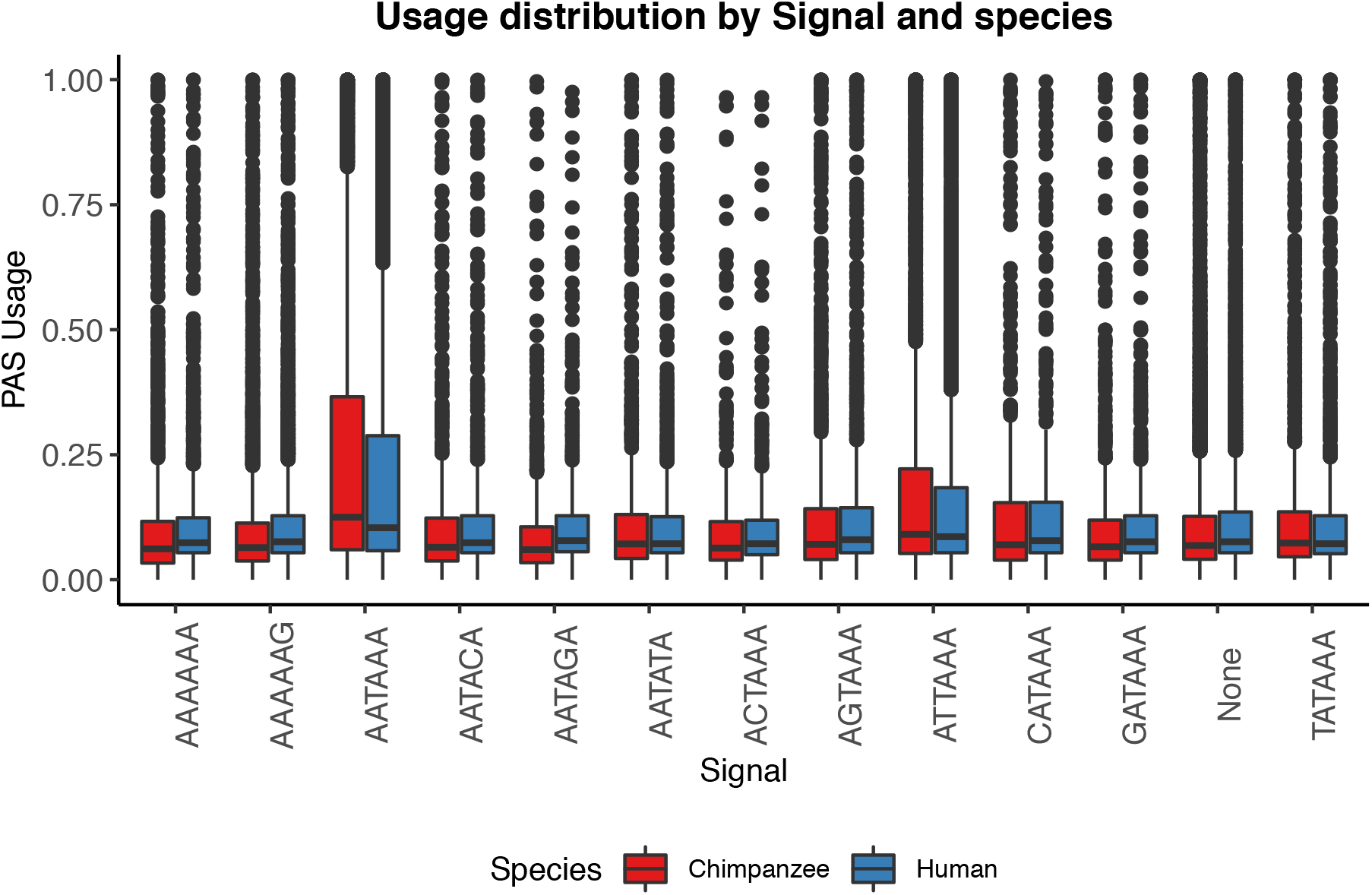
PAS with AATAAA and ATTAAA are used more often. Mean PAS usage of the top two signal site motifs in human and chimpanzee plotted by annotated signal site.

**Figure 1 - figure supplement 10:**
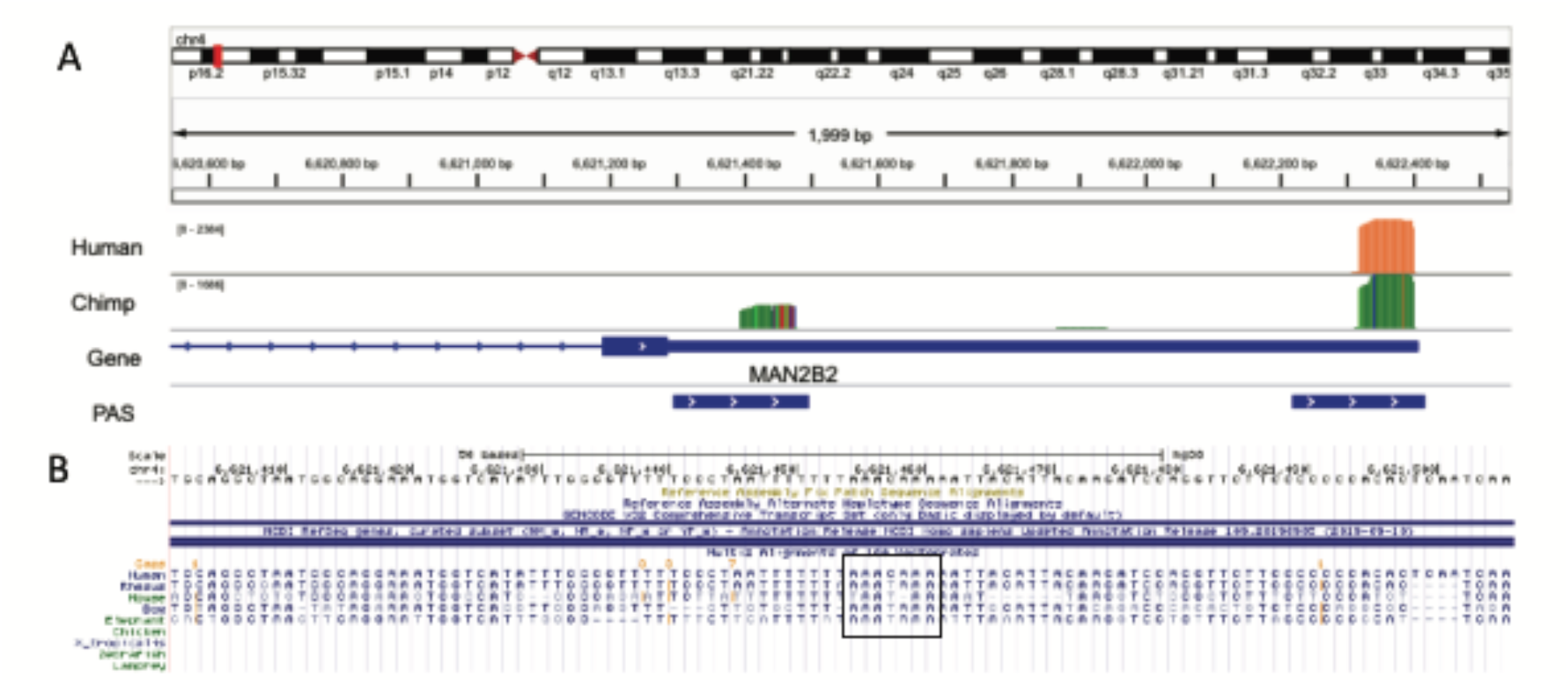
Chimp specific PAS likely due to loss of signal site in human lineage. A. IGV track for example of a chimpanzee specific PAS in MAN2B2 gene. Top track is merged coverage from 5 human nuclear 3’ seq libraries. Chimp track is merged coverage from 6 chimpanzee nuclear 3’ seq libraries lifted to human genome with CrossMap B. Sequence alignment for region upstream of proximal PAS from UCSC genome browser Black box indicates the signal site location. Canonical signal site is the ancestral state and was lost in the human lineage.

**Figure 1 - figure supplement 11:**
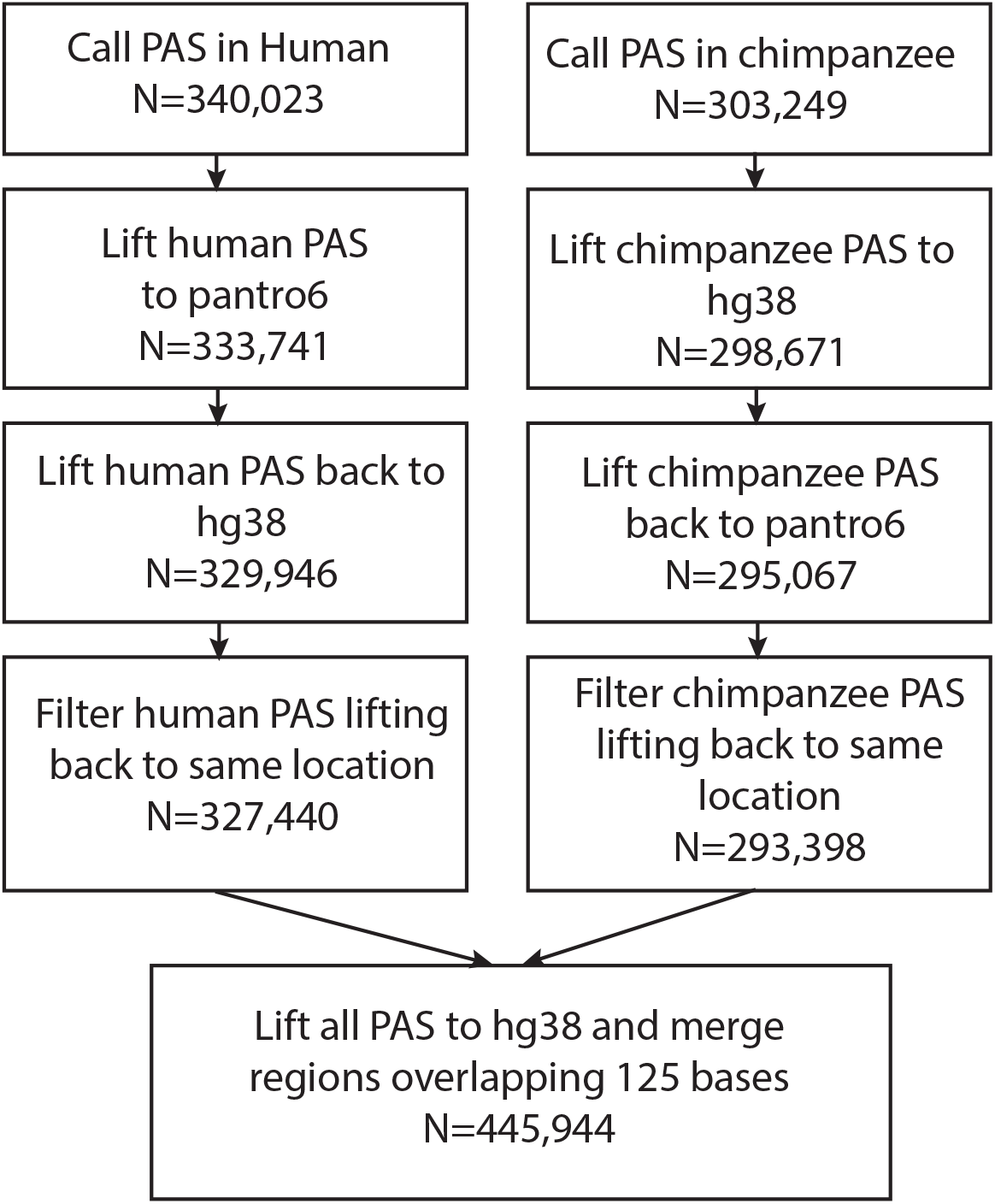
Reciprocal liftover pipeline. Reciprocal liftover pipeline for unfiltered PAS including the number of sites remaining at each step. Liftover using UCSC liftover tool and chain files downloaded from UCSC genome browser.

**Figure 2 - figure supplement 1:**
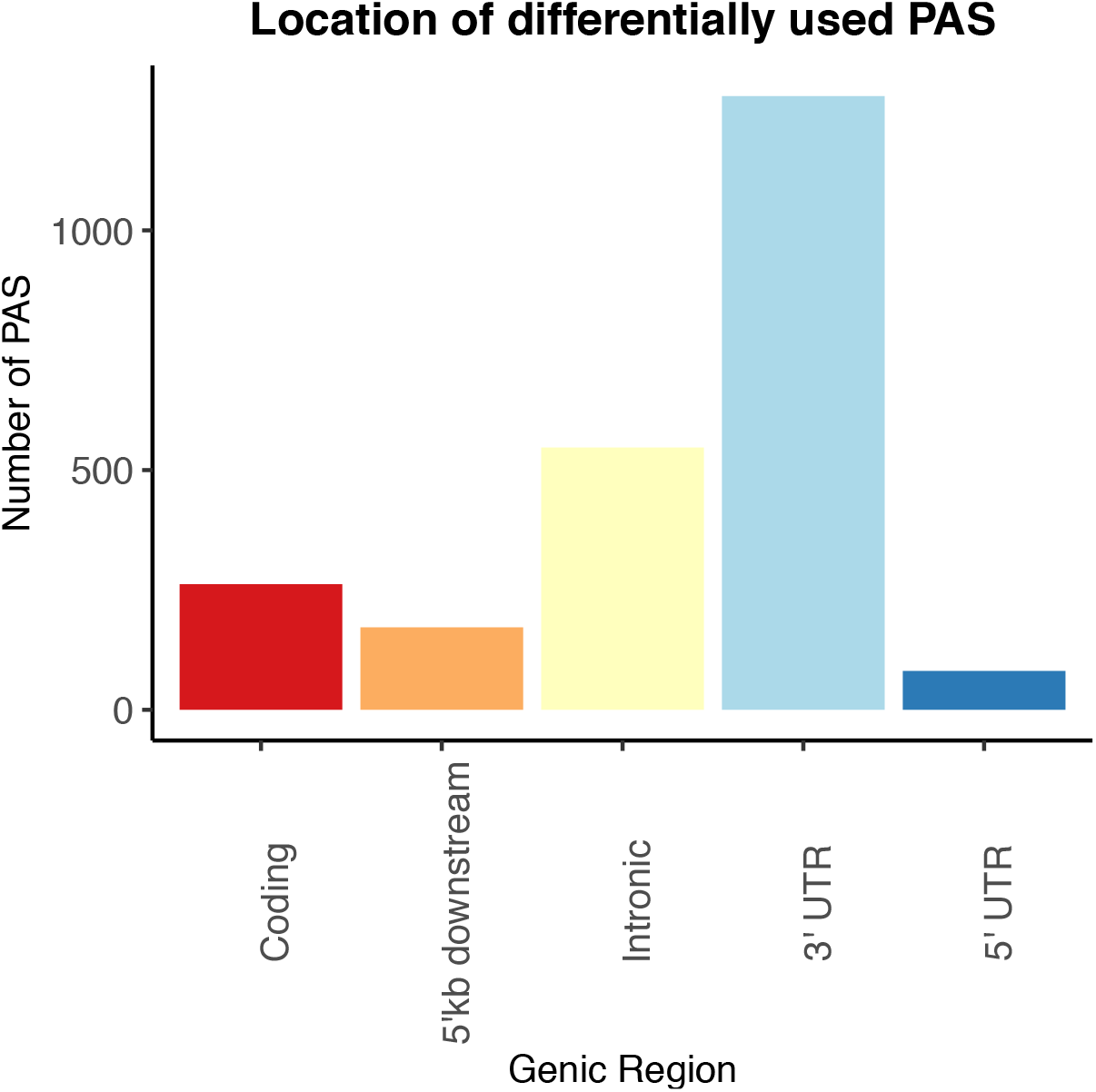
Genic Location of PAS differentially used between human and chimpanzee. Differentially used PAS (5% FDR) between human chimpanzee by genic annotation.

**Figure 2 - figure supplement 2:**
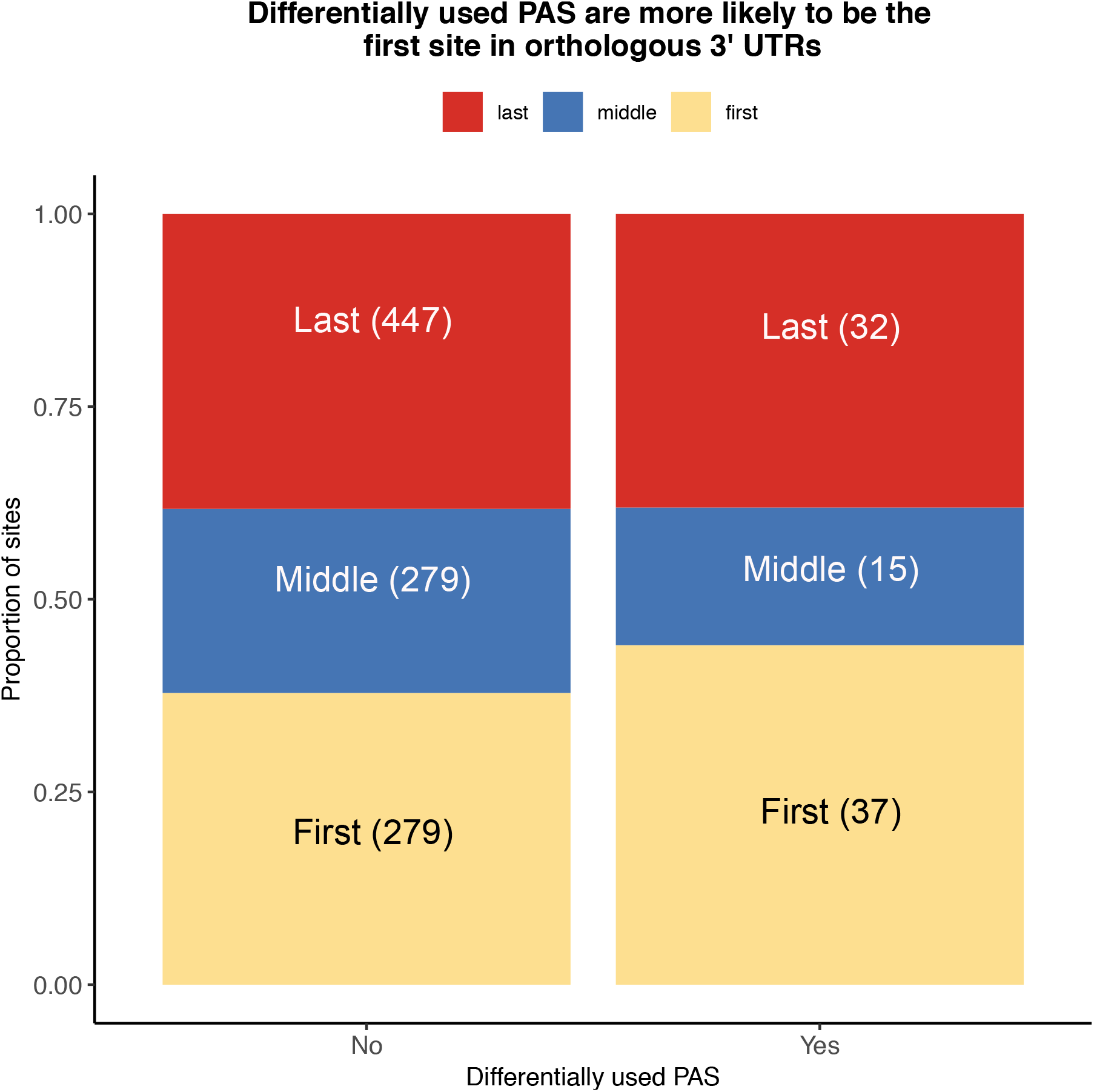
Location of PAS within Orthologous 3’ UTRs. Proportion of sites differentially used or conserved by whether they are the first (yellow), middle (blue), or last PAS (red) in orthologous exons.

**Figure 2 - figure supplement 3:**
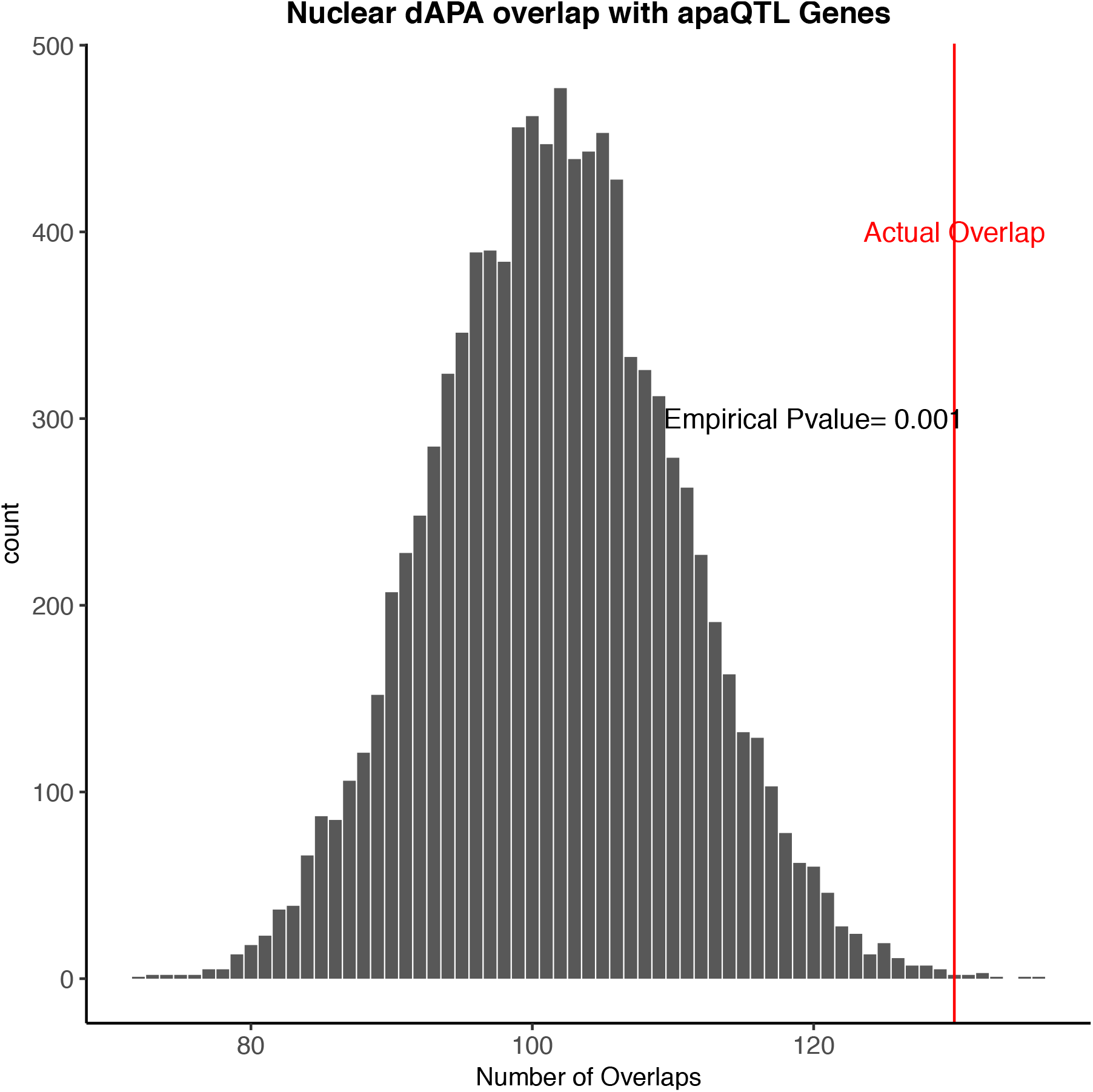
Genes with differentially used PAS are enriched for genes with apaQTL. 10,000 random subsamples of genes tested for differential APA and overlap with genes with apaQTLs Mittleman *et al*. Red line represents the actual overlap between genes with differential usage of at least one PAS and apaQTL genes.

**Figure 2 - figure supplement 4:**
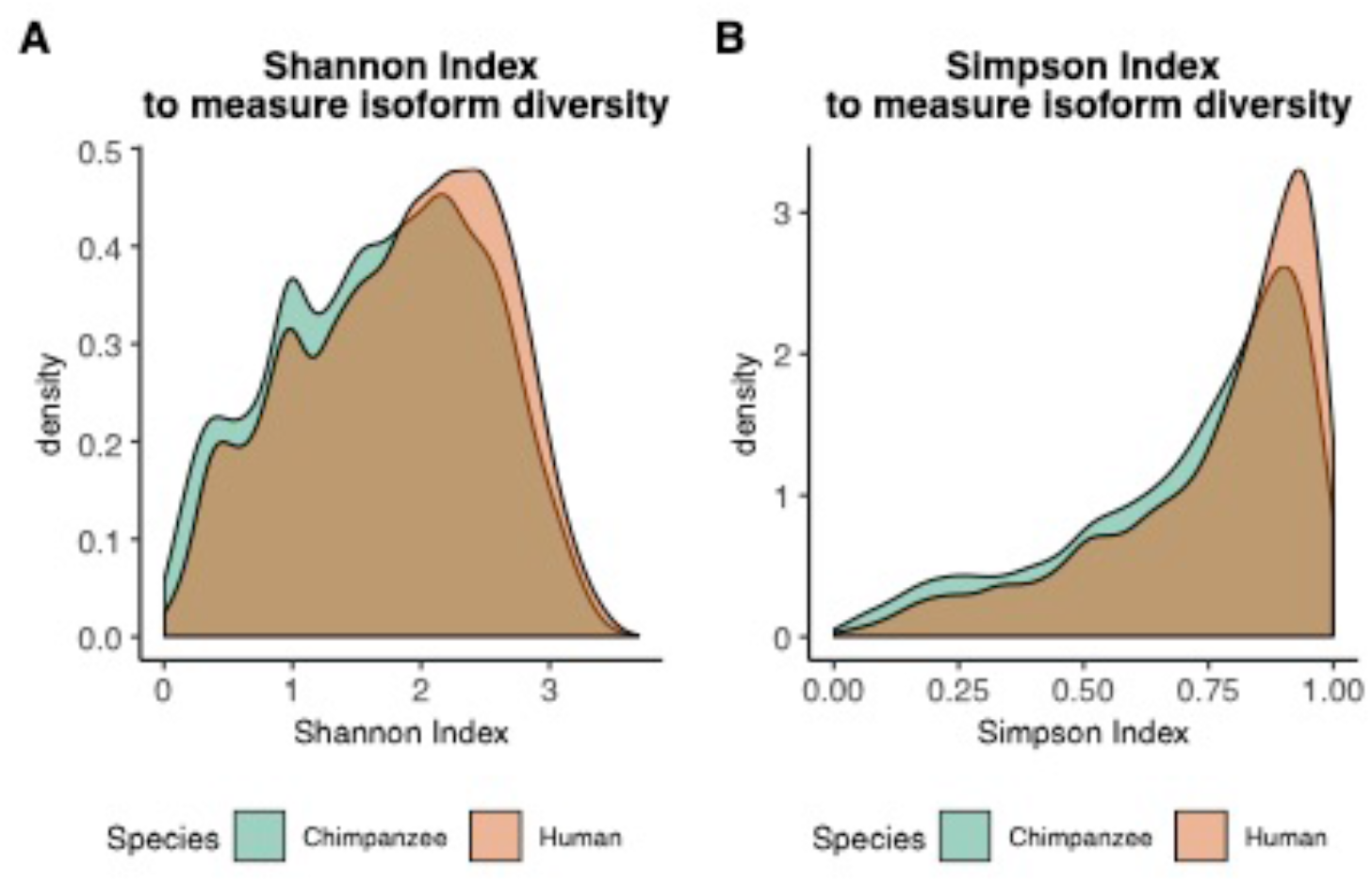
Information content measurement densities. A. Density of Shannon indices for all tested genes in human and chimpanzee 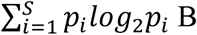. Density of Simpson indices for all tested genes in human and chimpanzee 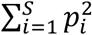

**Figure 2 - figure supplement 5:**
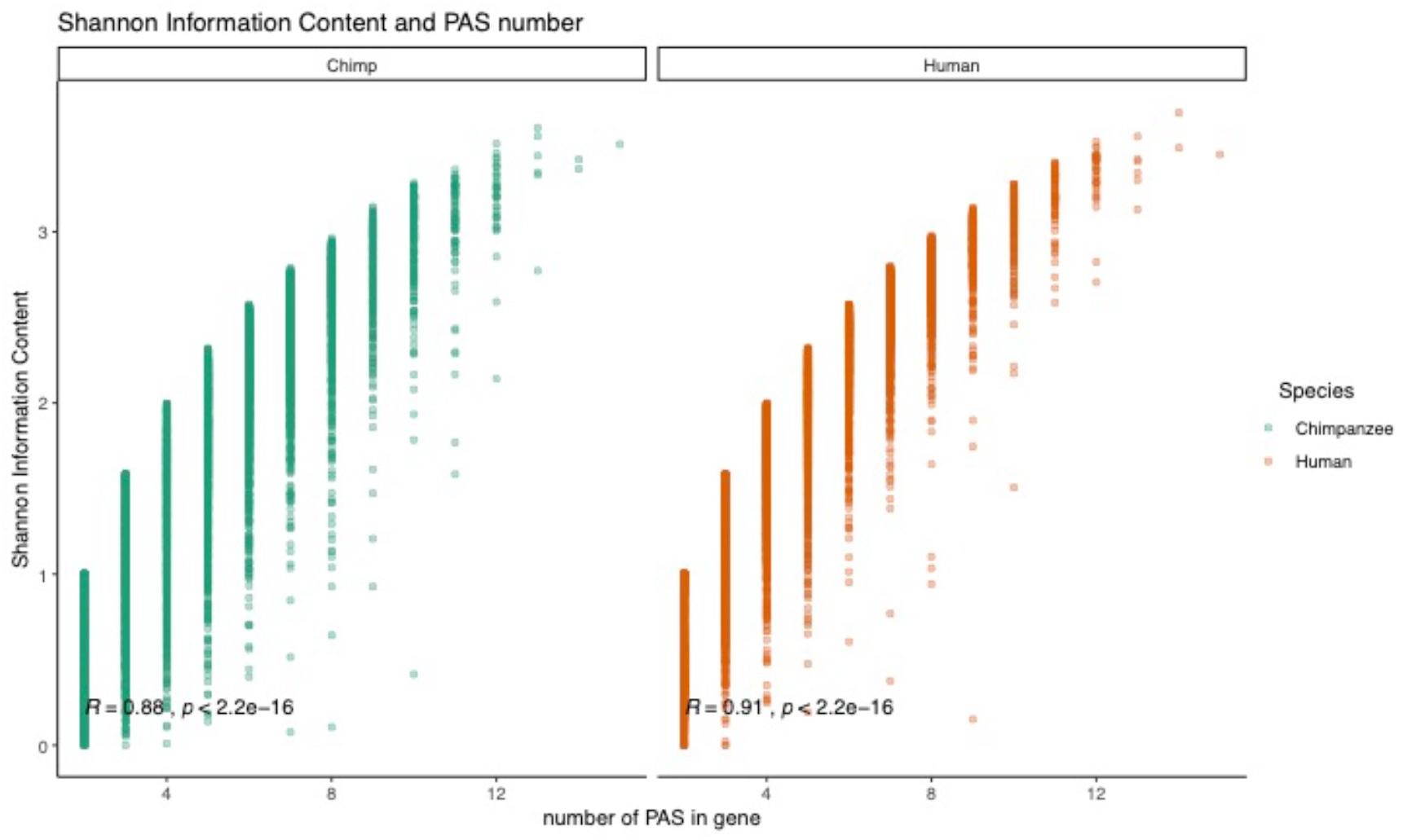
Relationship between Shannon index and PAS number. Shannon information index plotted against the number of PAS detect for each gene. Pearson’s correlation and significance in black

**Figure 2 - figure supplement 6:**
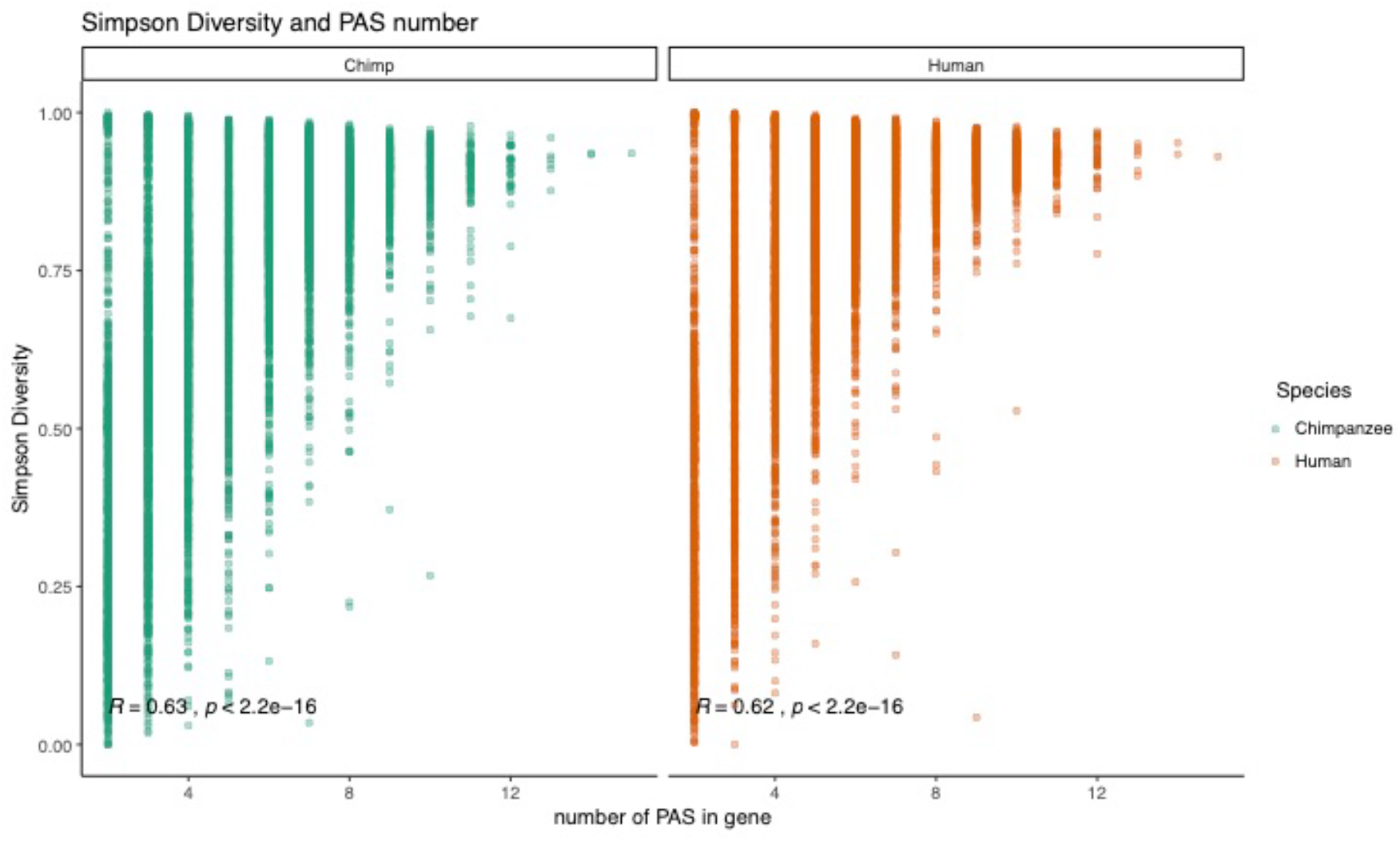
Relationship between Simpson diversity index and PAS number. Simpson’s diversity index plotted against the number of PAS detect for each gene. Pearson’s correlation and significance in black.

**Figure 2 - figure supplement 7:**
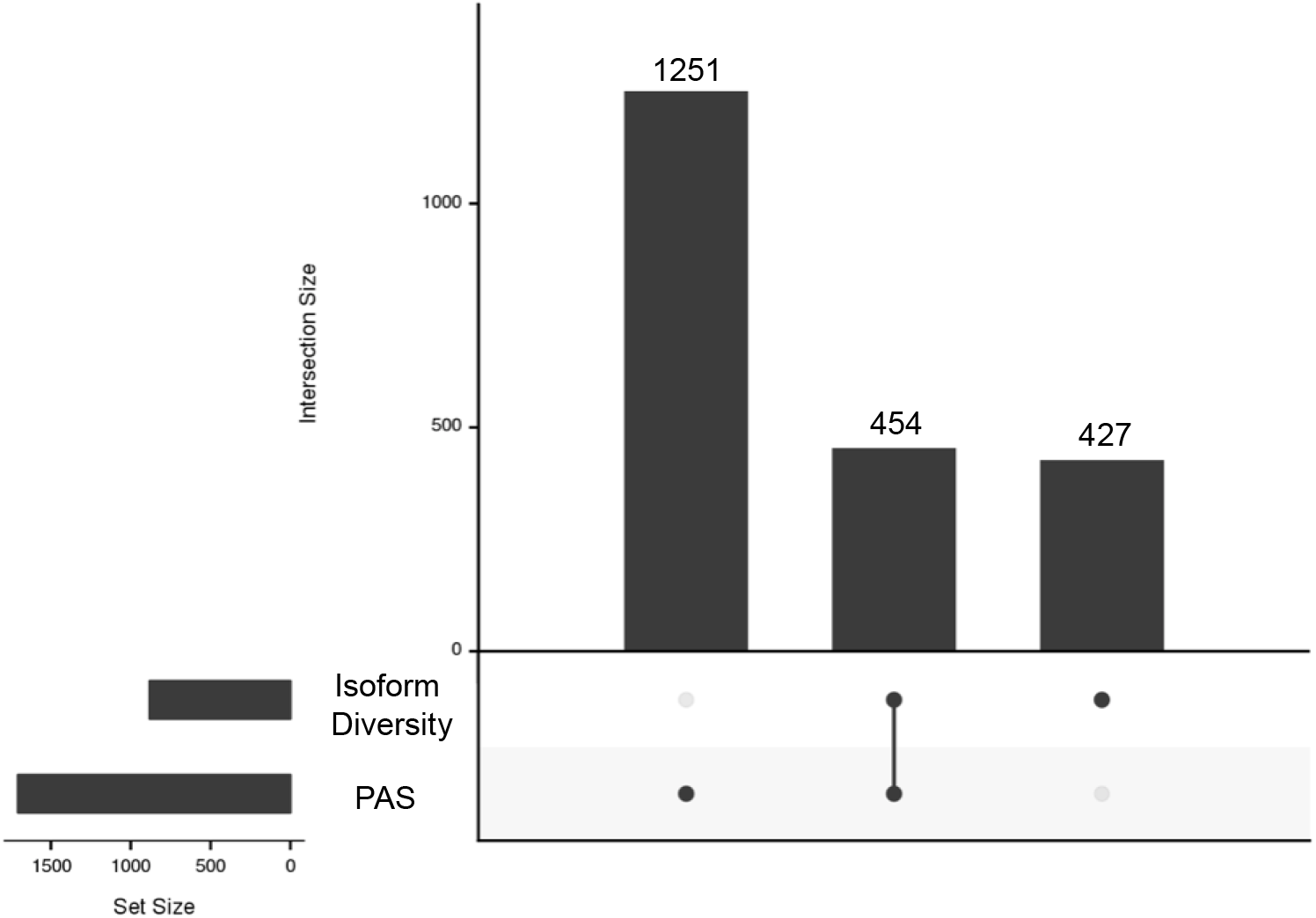
Intersection between genes with PAS and isoform diversity differences. 1251 genes have significant differences in PAS usage at between human and chimpanzee (left). 454 genes have significant differences in APA between humans and chimpanzee in PAS usage and in isoform diversity (middle). 427 genes with differences in isoform diversity level only (right).

**Figure 2 - figure supplement 8:**
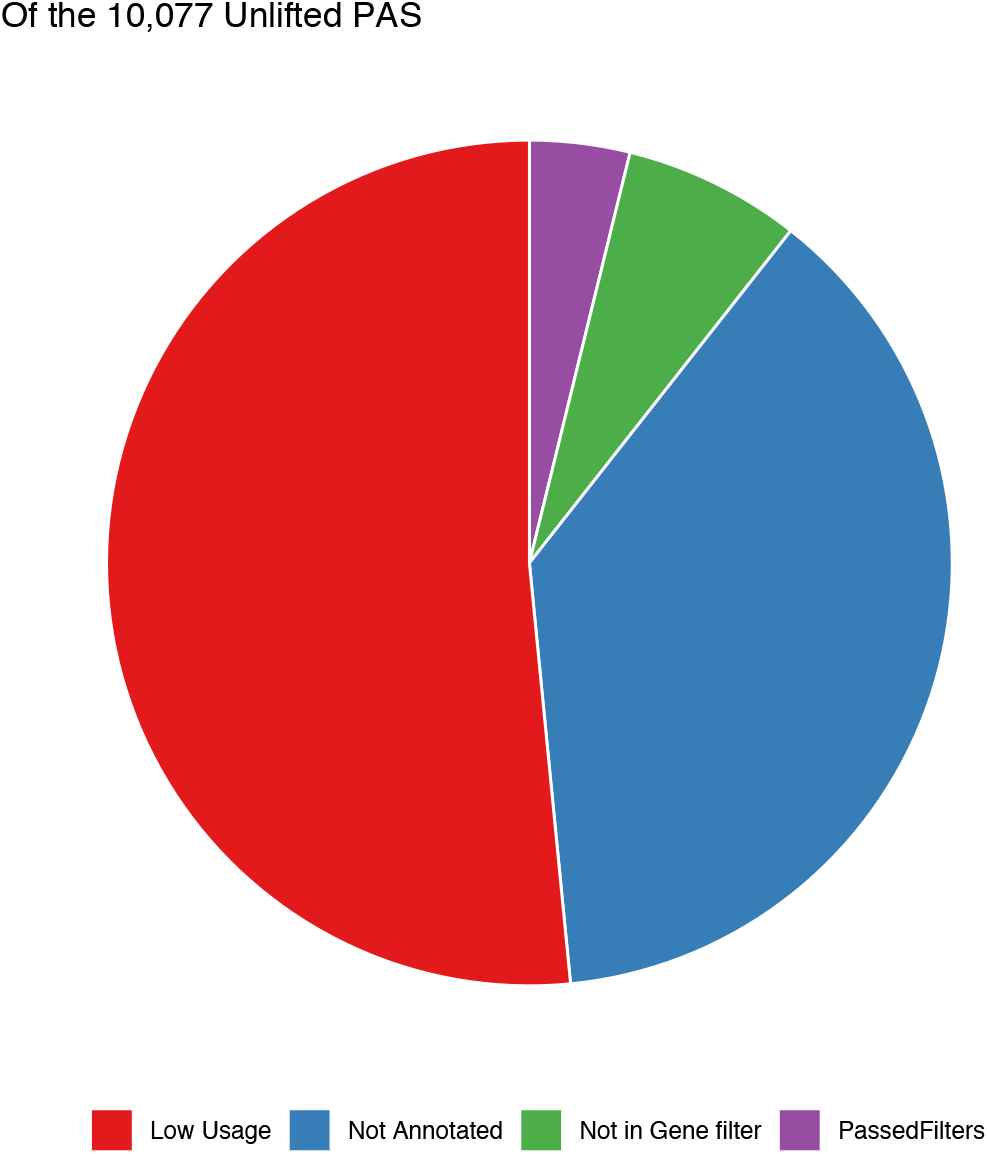
PAS that do not lift from human to chimp. Of the 10,077 PAS that do not reciprocally lift from human to chimp, distribution of where sites are filtered out. Most are lost due to not mapping to genes or due to low usage (likely noise)

**Figure 3 - figure supplement 1:**
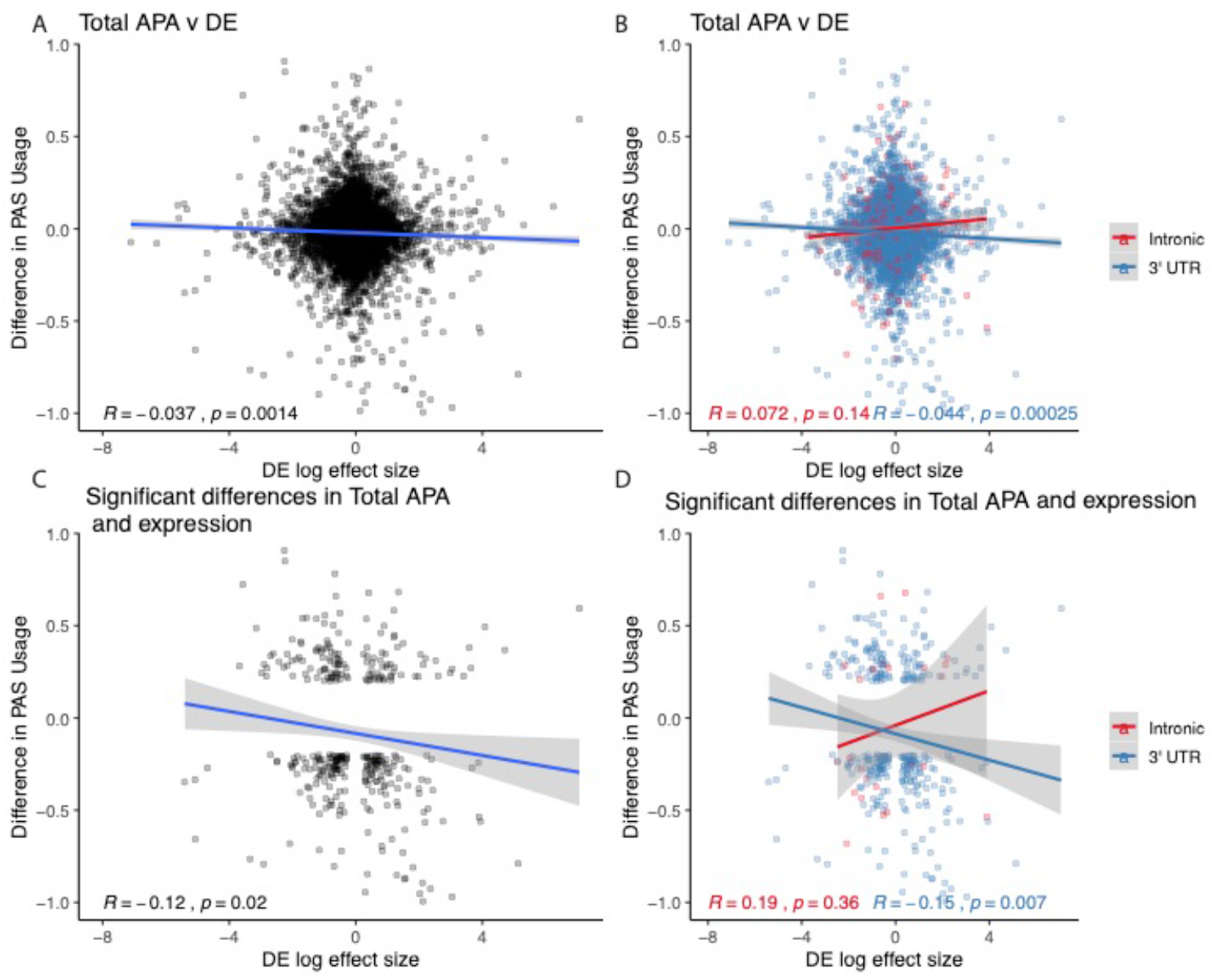
Figure 3 relationships expanded to total usage. A. Total mRNA ΔPAU for top intronic or 3’ UTR PAS per gene plotted against differential effect size from differential expression analysis. B. Total mRNA ΔPAU for top intronic or 3’ UTR PAS per gene plotted against differential effect size from differential expression analysis for genes with significant differences in each phenotype at 5% FDR. C. Total mRNA ΔPAU for top intronic or 3’ UTR PAS per gene plotted against differential effect size from differential expression analysis. D. Total mRNA ΔPAU for top intronic or 3’ UTR PAS per gene plotted against differential effect size from differential expression analysis for genes with significant differences in each phenotype at 5% FDR. In all panels, we calculated the linear regression and Pearson’s correlation with the r package ggpubr. In B and D, we colored the points and regression line by genic location. In all panels, negative ΔPAU and DE effect sizes represent upregulation in chimpanzees.

**Figure 3 - figure supplement 2:**
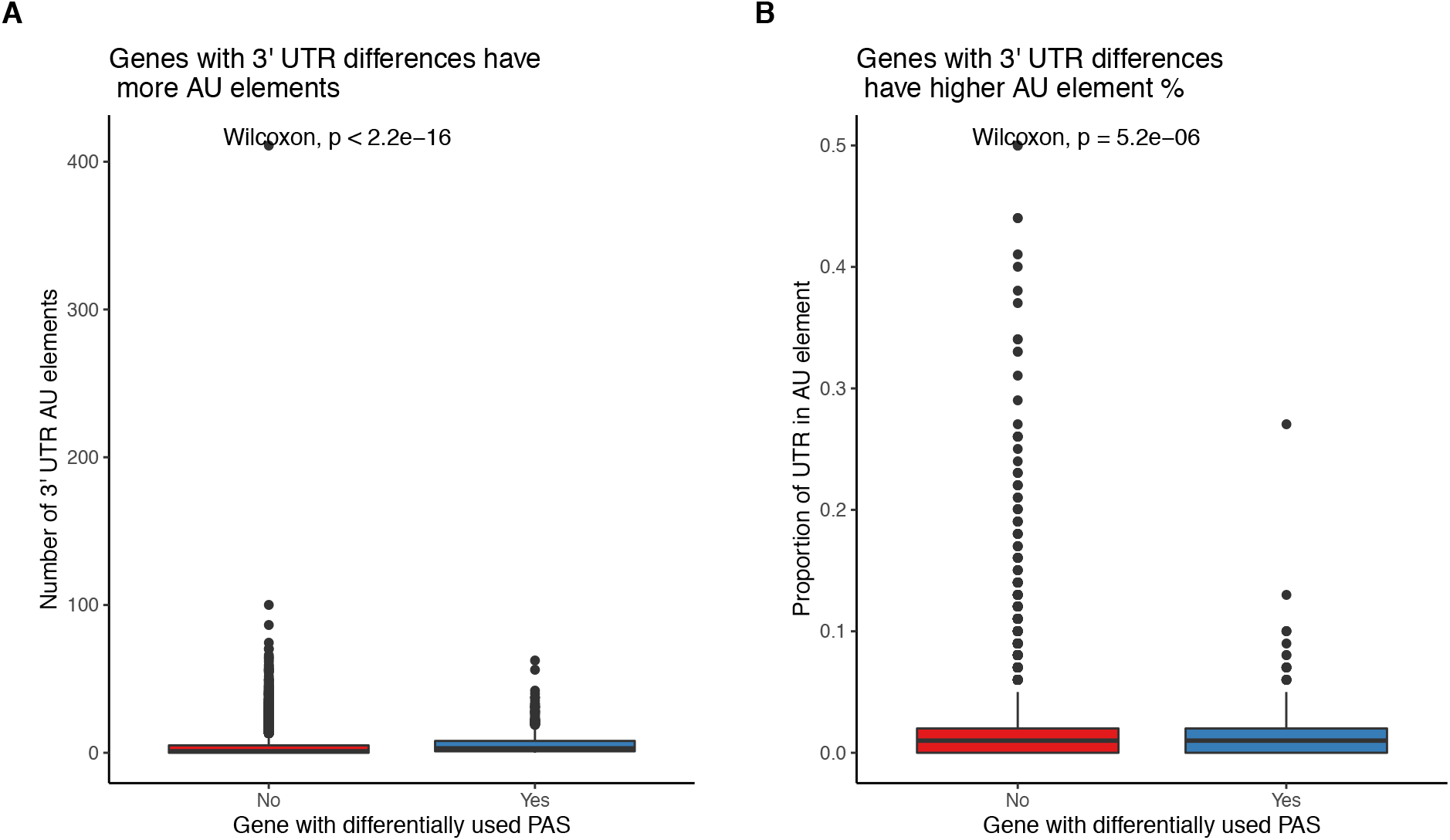
Differentially used 3’ UTR PAS have higher AU element content. A. Genes with at least one differentially used 3’ UTR PAS have more AU elements in their 3’ UTRs. B. Genes with at least one differentially used 3’ UTR PAS have a higher proportion of their 3’ UTRs covered by AU elements.

**Figure 3 - figure supplement 3:**
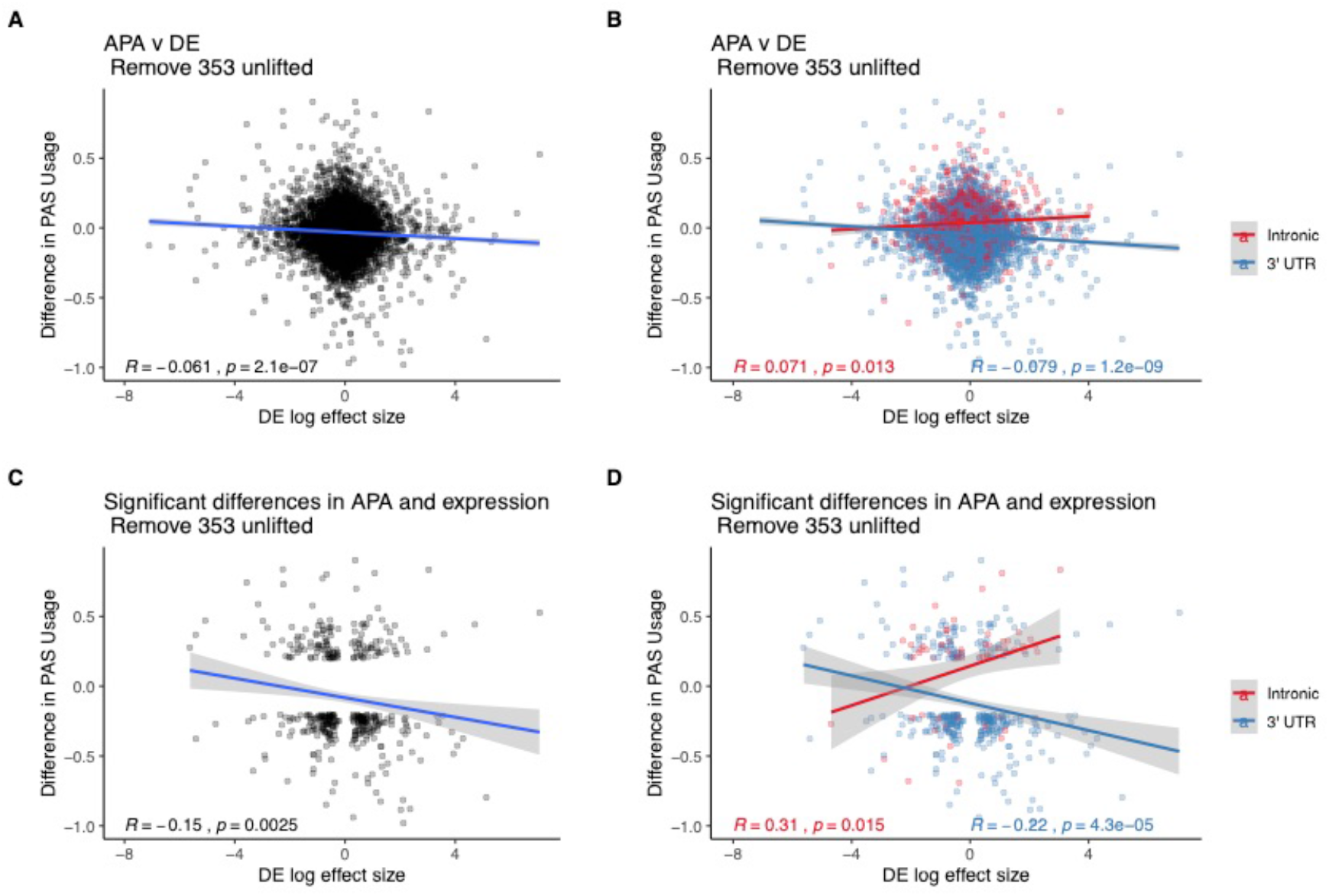
Figure 3 without genes affected by liftover. A. ΔPAU for top intronic or 3’ UTR PAS per gene plotted against differential effect size from differential expression analysis. B. ΔPAU for top intronic or 3’ UTR PAS per gene plotted against differential effect size from differential expression analysis for genes with significant differences in each phenotype at 5\% FDR. C. ΔPAU for top intronic or 3’ UTR PAS per gene plotted against differential effect size from differential expression analysis. D. ΔPAU for top intronic or 3’ UTR PAS per gene plotted against differential effect size from differential expression analysis for genes with significant differences in each phenotype at 5% FDR. In all panels, we calculated the linear regression and Pearson’s correlation with the r package ggpubr. In B and D, we colored the points and regression line by genic location. In all panels, negative ΔPAU and DE effect sizes represent upregulation in chimpanzees.

**Figure 3 - figure supplement 4:**
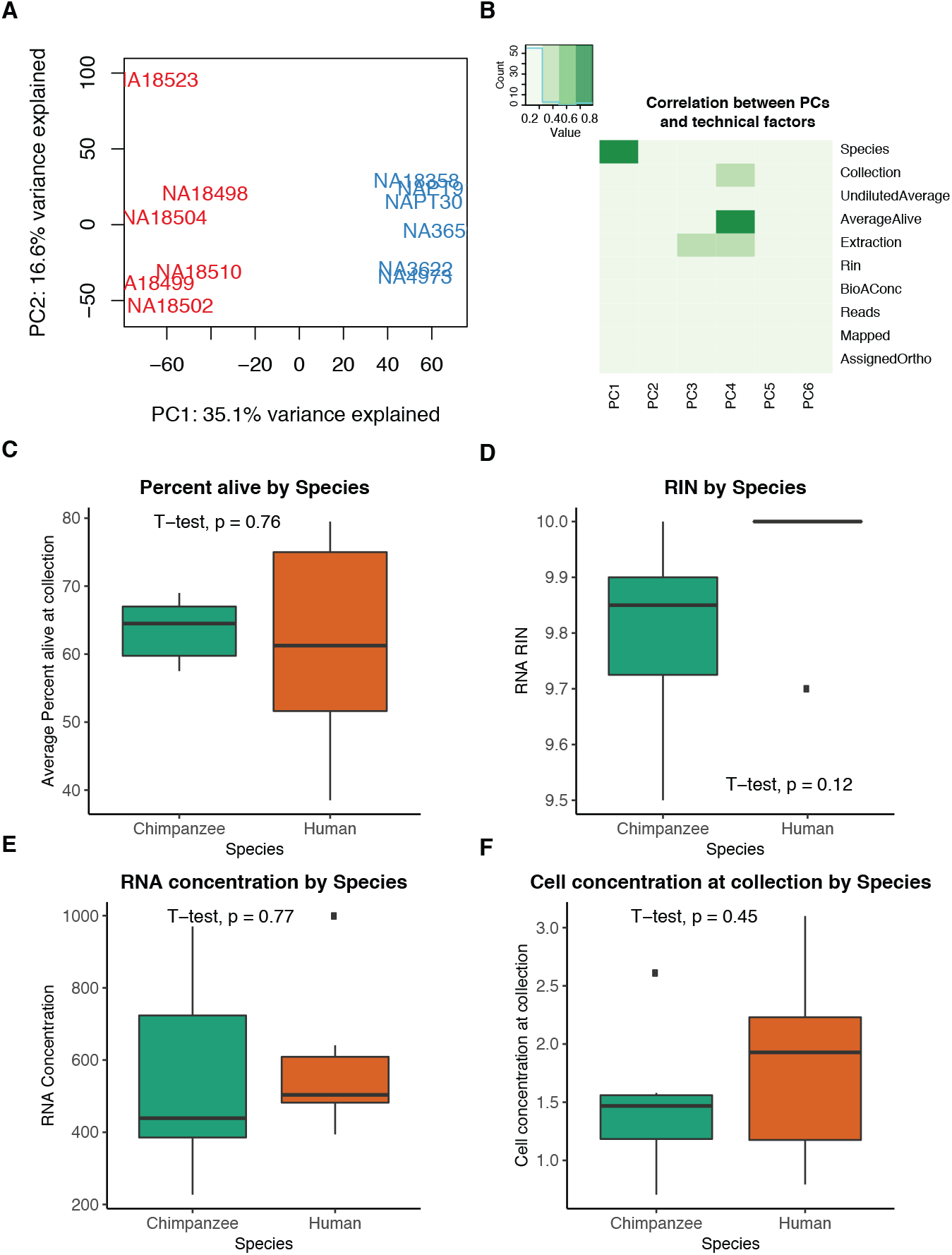
Differential expression quality control plots. A. First two principal components (PCs) in gene expression variation. B. Heatmap representing correlation between technical factors and PCs. Explanation of Y axis factors and values available in Supplementary Table 3 C. Percent of live cells as calculated by trypan blue staining at collection is not confounded by species. D. RIN scores reported by bioanalyzer at RNA-seq library generation are not confounded by species. E. RNA concentrations reported by bioanalyzer at RNA-seq library generation are not confounded by species. F. Cell concentrations at time of collection are not confounded by species.

**Figure 4 - figure supplement 1:**
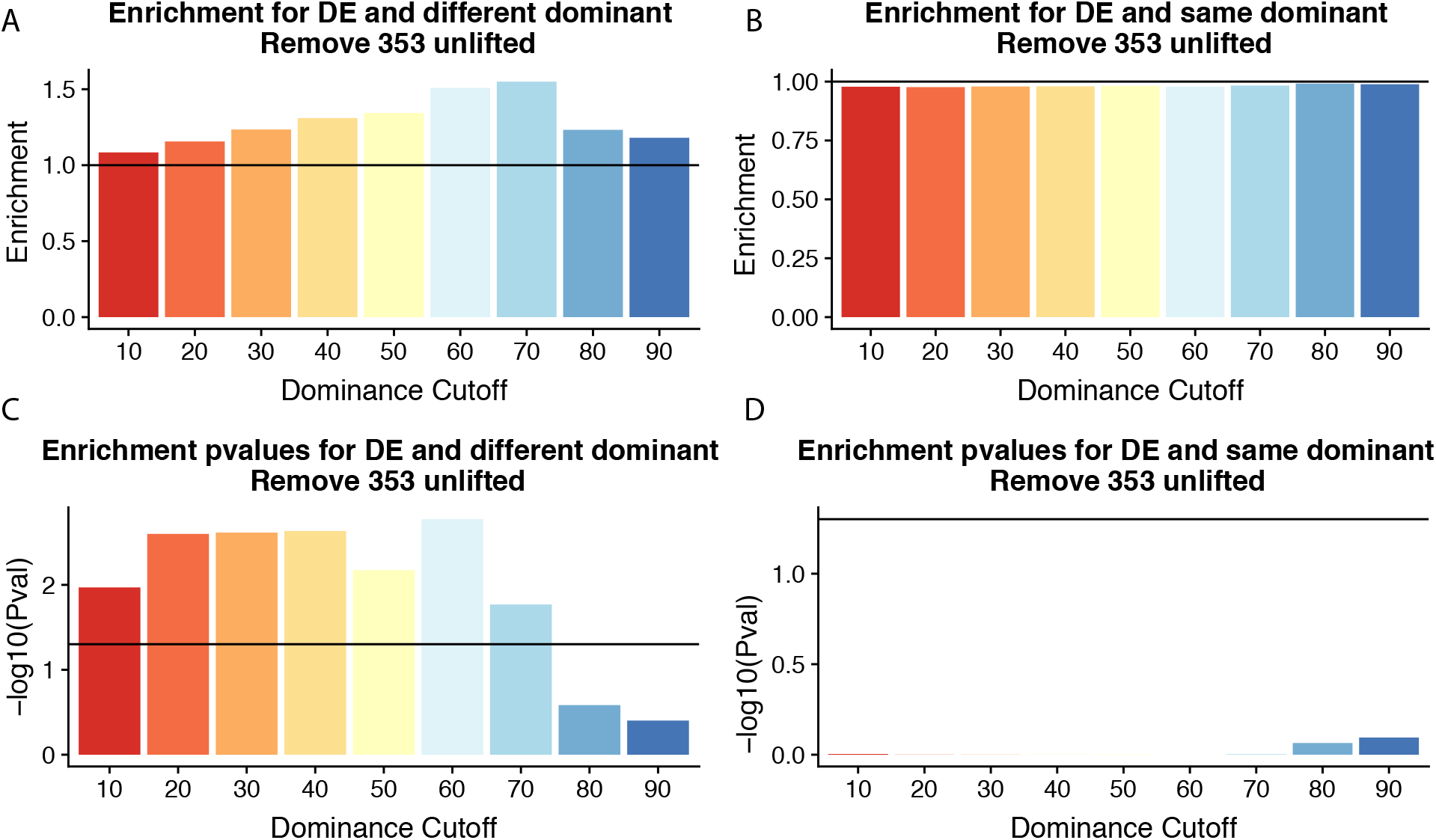
Figure 4 without genes affected by liftover. A. Enrichment of genes with the different (left) or same (right) dominant PAS by dominant cutoff in differentially expressed genes after removing genes likely affected by liftover. B. −log_10_(p-values) for enrichments in A calculated with hypergeometric tests. Horizontal line represents p= 0.05.

**Figure 5 - figure supplement 1:**
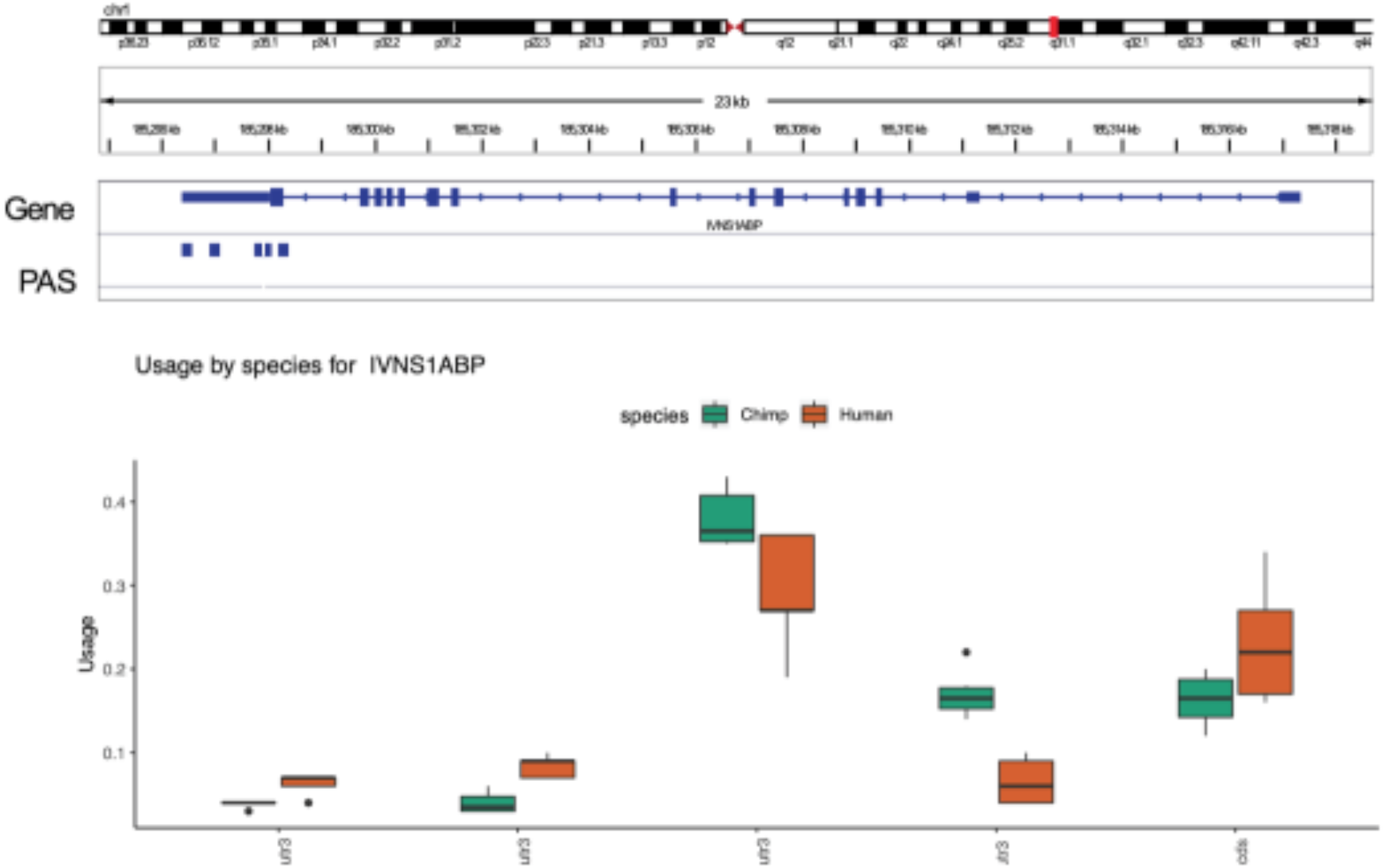
Gene with significant differences in isoform diversity only. Human and chimpanzee usage for 5 PAS identified in the IVNS1ABP gene. None of the PAS measured have significant differences in usage at 5% FDR. IVNS1ABP is not differentially expressed.

**Figure 5 - figure supplement 2:**
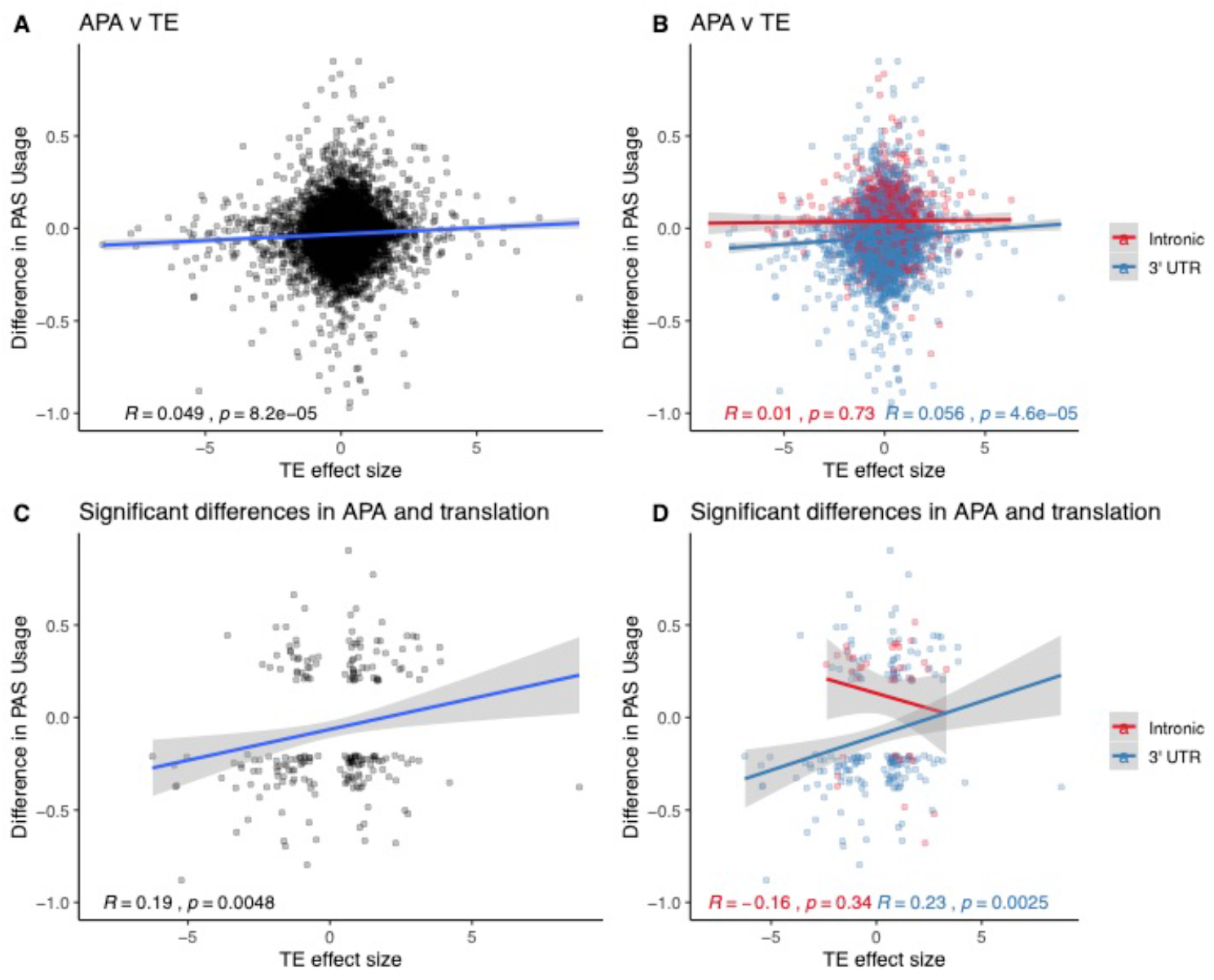
Relationship between ΔPAU and differential translation effect sizes. A. ΔPAU for top 3’ UTR and intronic PAS plotted against differential translation (TE) effect size as reported by Wang et al. B. ΔPAU for top 3’ UTR and intronic PAS plotted against TE effect size as reported by Wang et al. separated by genic location. C. ΔPAU for top 3’ UTR and intronic PAS with significant differences in usage plotted against TE effect size for significant genes (5% FWER) as reported by Wang et al. D. ΔPAU for top 3’ UTR and intronic PAS with significant differences in usage plotted against TE effect size for significant genes (5% FWER) as reported by Wang et al. separated by genic location. Linear regression line was plotted and Pearson’s correlation was calculated for data in each panel.

**Figure 5 - figure supplement 3:**
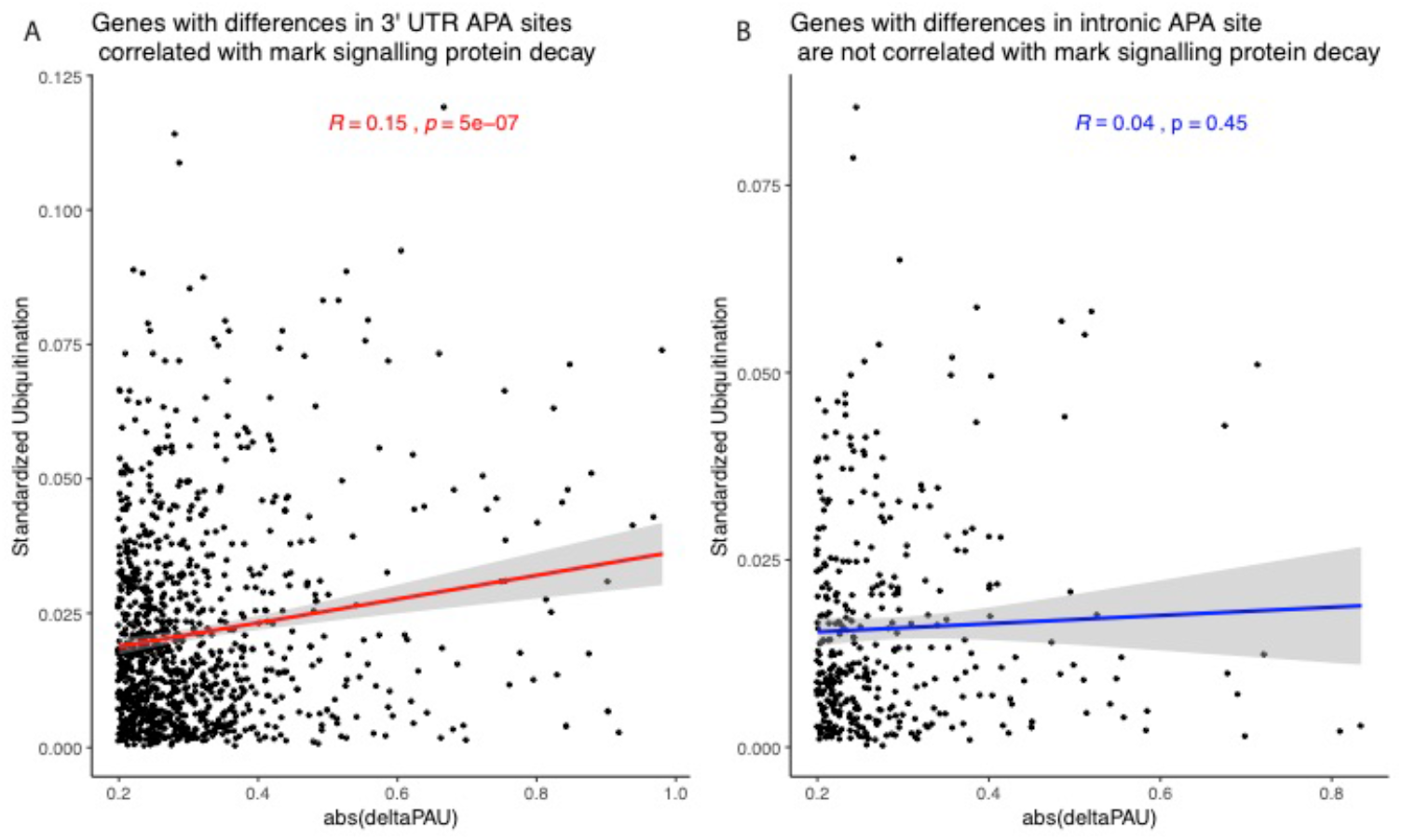
Relationship between APA differences and protein decay mark. A. Absolute value of ΔPAU for 3’ UTR PAS with significant difference at site level plotted against the number of ubiquitination marks in the gene standardized by the number of amino acids. Regression line and Pearson’s correlation are plotted in red. B. Absolute value of ΔPAU for intronic PAS with significant difference at site level plotted against the number of ubiquitination marks in the gene standardized by the number of amino acids. Regression line and Pearson’s correlation are plotted in blue.

**Figure 5 - figure supplement 4:**
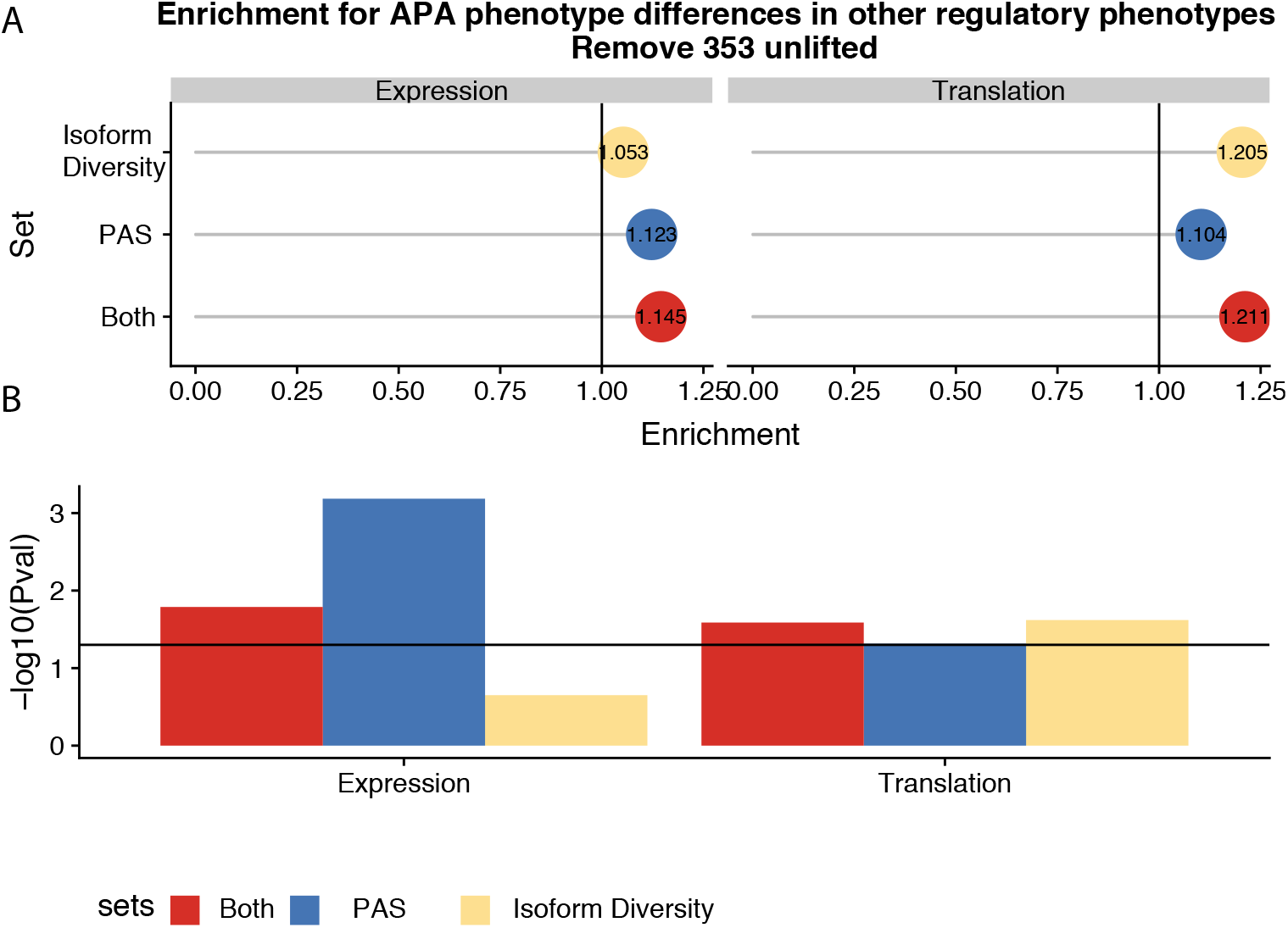
Figure 5 without genes affected by liftover. A. Enrichment of genes with differences in isoform diversity, PAS usage, or both within differential expressed genes and differentially translated genes after removing genes likely affected by liftover. Differentially translated genes reported by Wang et al. B. −log_10_(p-values) for enrichments in A calculated with hypergeometric tests. Horizontal line represents p= 0.05.

**Figure 6 - figure supplement 1:**
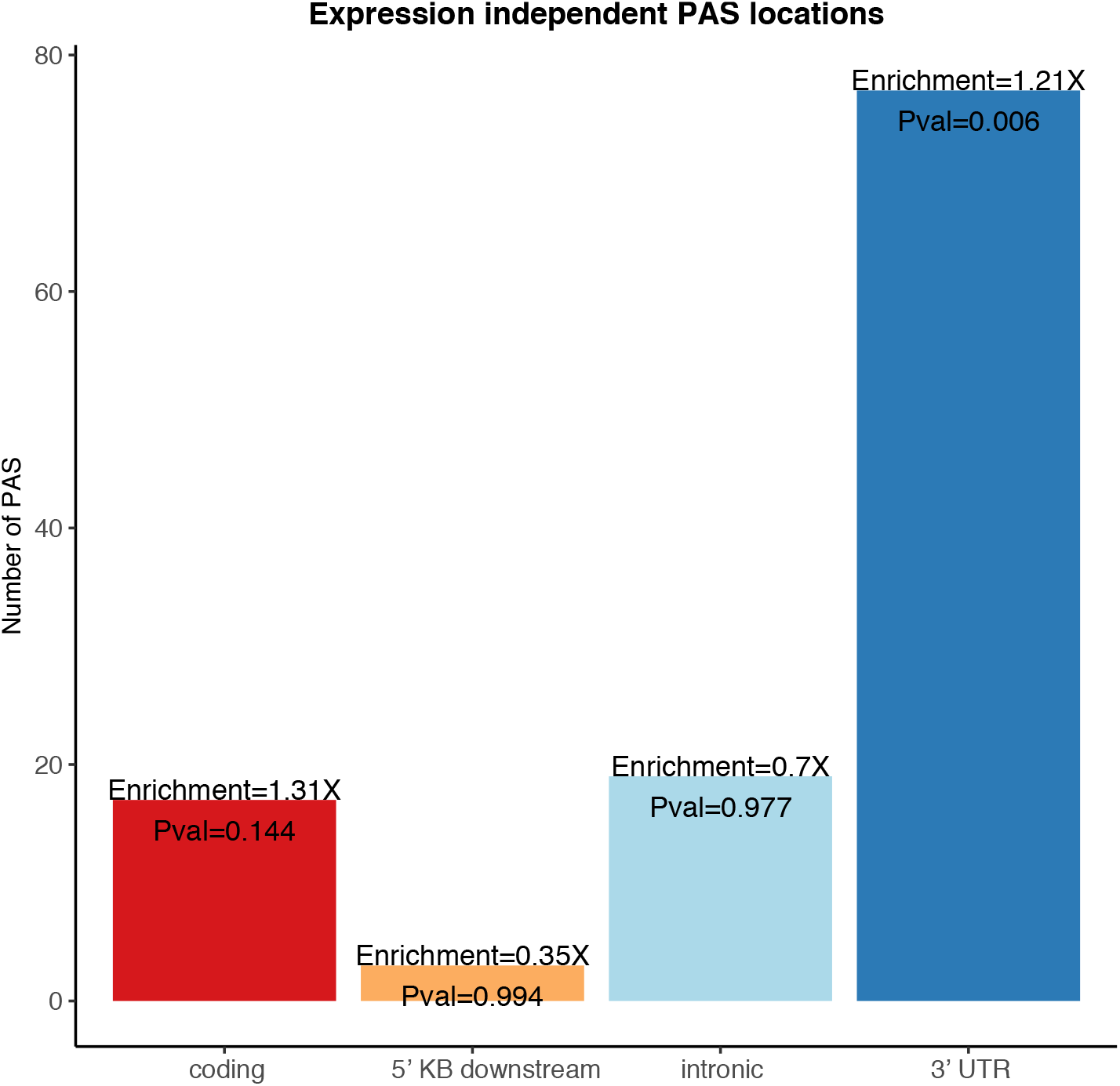
Enrichment for 3’ UTR PAS in genes differentially expressed in protein and not in mRNA. Genic location enrichments for the PAS in genes differentially expressed at protein level but not mRNA level among all differentially used PAS. P-values were calculated with a hypergeometric test.

**Figure 6 - figure supplement 2:**
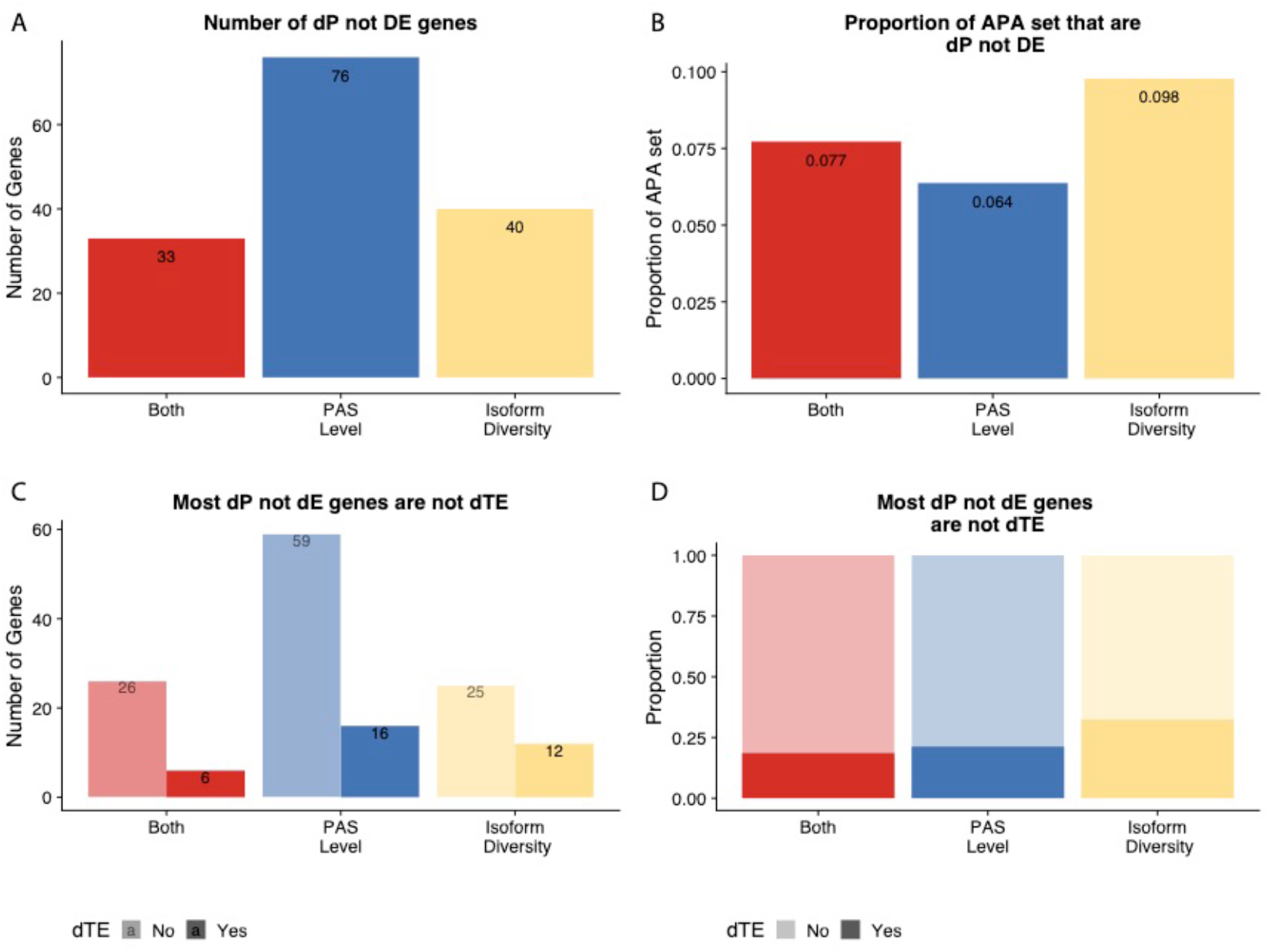
Figure 6 without genes affected by liftover. A. Number of genes with differences in isoform diversity, PAS usage or both differentially expressed in protein (5% FDR) but not in mRNA (5% FDR). Genes differentially expressed in protein from Khan et al. B. Proportion of genes with differential isoform diversity, PAS usage or both that are differentially expressed in protein (5% FDR), but not mRNA (5% FDR). C. Genes reported in separated by genes differentially translated at 5% FDR. Differentially translated gene reported in Wang et al. D. Genes differentially expressed in protein but not in mRNA, colored by differences in APA. Proportion of genes in the set differentially translated at 5% FDR.

## Supplemental Tables

**Supplemental Table 1: PAS Differential Usage results.** Column names as described-PAS: polyadenylation site, gene: gene (cluster in leafcutter), PAS_logeffectsize: log effect size for differential usage of the PAS between human and chimpanzee, PAS_deltaPAU: difference in polyadenylation site usage between human and chimpanzee (delta PSI in leafcutter), Gene_logLR: log likelihood ratio for differential usage of any PAS in the gene. (cluster likelihood ratio in leafcutter), Gene_adjustedPvalue: adjusted pvalue for differential usage of the any PAS in the gene (cluster pvalue in leafcutter)

**Supplemental Table 2: 3’ Sequencing metadata** Column names as described-Species: Cell line species, Lines: Cell line ID, Fraction: Cellular fraction, CollectionDate: Date of cell harvest and nuclear isolation, Extraction_date: Date of RNA extraction, Collection_person: Author initial for who processed cell harvest and nuclear isolation, UndilutedAverage: Average of 2 cell count measurements 1×10^6^, AverageAlive: Average of 2 cell live dead counts - calculated with trypan blue stain, Concentration: Extracted RNA concentration (ng/ml), RIN: RIN score for extracted RNA, 260.280.Ratio: 260/280 ratio calculated on nanodrop, Library: 3’ Seq library date, Reads: Number of sequenced reads, Mapped_wMP: Number of Mapped reads before removing reads likely due to misprimming, Mapped_Clean: Number of Mapped reads after removing reads likely due to misprimming

**Supplemental Table 3: Metadata for RNA sequencing data** Column names as described-Species: Cell line species, Lines: Cell line ID, Collection_person: Author initial for who processed cell harvest and nuclear isolation, UndilutedAverage: Average of 2 cell count measurements 1×10^6^, AverageAlive: Average of 2 cell live dead counts - calculated with trypan blue stain, CollectionDate: Date of cell harvest and nuclear isolation, Extraction: Date of RNA extraction, RIN: RIN score for extracted RNA, BioAConc: RNA concentration (ng/ul), Reads: Number of Sequenced reads, Mapped: Number of mapped reads, AssignedOrtho: Number of mapped reads assigned to orthologous exons.

**Supplemental Table 4: Differential Expression Results** Column names as described-Differential expression results from limma. gene: tested gene, logFC: log 2 fold change in normalized gene expression, adj.P.Val: BH adjusted pvalue from t test, B: Beta value, t: t statistic

